# Response dynamics of discrete subiculum→retrosplenial cortex projections underlying trace fear conditioning

**DOI:** 10.1101/2025.06.12.659393

**Authors:** Thomas E. Bassett, Yi-Zhi Wang, Elizabeth M. Wood, Hui Zhang, Zorica Petrovic, Vivien Prifti, Vladimir Jovasevic, Naoki Yamawaki, Lynn Ren, Natalia Khalatyan, Viktoriya Grayson, Jeffrey N Savas, Jelena Radulovic, Ana Cicvaric

## Abstract

Associating events separated in time depends on the CA1, subiculum (SUB), and retrosplenial cortex (RSP). The degree to which their connectivity and underlying circuit mechanisms contribute to the association of such temporally discontiguous events is not known. Here we showed, using trace fear conditioning (TFC), wherein mice learn to associate tone and shock separated by a temporal trace, that molecularly distinct excitatory VGluT1^+^ and VGluT2^+^ SUB→RSP projections subserve the associative and temporal components of TFC. During trace memory formation, VGluT2^+^ SUB→RSP projections showed increased and decreased bulk calcium activity at tone and trace onset, respectively, an activity pattern that was reestablished during memory recall. Such pattern was not observed in CA subfields, suggesting that associative and temporal components of TFC are integrated at the SUB or SUB→RSP synapses before being presented to the RSP. Our findings establish a circuit mechanism for representing complex temporal information in episodic memory.

## Introduction

Episodic memory is defined as a form of consciously recallable long-term memory encompassing details of personally experienced events, including their temporal, spatial, and relational properties.^1,2^ Beyond simply situating a memory in time, the temporal component of episodic memory holds information regarding the order in which events happened, their durations, and the time intervals between them.^3^ Collectively, these temporal properties allow individuals to integrate discrete events separated in time into a cohesive and meaningful memory.

The capacity to bind discontiguous events can be tested in rodent models with paradigms such as trace fear conditioning (TFC), in which animals learn to associate a tone and foot-shock separated by a temporal gap.^4–6^ TFC requires coordinated activity across multiple components of both hippocampal complex and cerebral cortex to support proper memory formation and recall.^4,7^ Inhibition of the dorsal hippocampal CA1 subfield significantly impairs both conditioning and recall in TFC.^8–10^ In addition, the entorhinal cortex contributes to TFC through NMDAR-mediated glutamatergic projections into the CA1,^4,11,12^ while inhibition of the dorsal CA1 projections to the dorsal subiculum (SUB) during the temporal gap impairs TFC memory formation.^13^ Finally, projections from the SUB back into the EC are necessary for TFC memory recall, thus forming a critical circuit within the hippocampal formation underpinning TFC.^10,13^ In the cortex, the retrosplenial cortex (RSP) contributes significantly to TFC memory formation and extinction,^14–16^ and plays a key role in episodic memory as a temporal processing hub.^17^ However, while the individual functions of the hippocampal formation and the RSP are well-established, if and how these regions communicate during TFC remains unexplored.

Our previous work demonstrates that the hippocampal complex interfaces with the RSP via inhibitory projection from the CA1^18^ and glutamatergic projections originating from the SUB, comprising of vesicular glutamate transporter 1 (VGluT1^+^) and vesicular glutamate transporter 2 (VGluT2^+^) positive pyramidal neurons,^19,20^ with different contributions to the formation of ssociative context memories.^18,19^ Using TFC in combination with chemogenetic inhibition, we found that while SUB→RSP VGluT1^+^ and VGluT2^+^ projections redundantly encoded the aversive tone-shock association, only VGluT2^+^ projections were required for the processing of the temporal trace. Using fiber photometry, we showed that SUB→RSP VGluT2^+^ afferents exhibited biphasic calcium transients, with an early calcium peak occurring at tone onset, followed by a pronounced dip during the trace period. This pattern emerged during training, reappeared during recall, and was absent in the pseudoconditioned control group, demonstrating that VGluT2^+^ SUB→RSP activity patterns specifically encoded the temporal gap. The observed activity pattern was not found in the CA subfields.

## Results

### VGluT2^+^ SUB→RSP afferents are specifically required for processing a temporal trace separating a cue and a shock

In delay fear conditioning (DFC), mice are exposed to a presentation of a tone terminating with footshock, whereas trace fear conditioning (TFC) introduces a temporal gap (trace) between them. In TFC -unlike DFC-the dorsal hippocampus (DH) ^21^ and RSP^22^ are necessary for the association of tone and shock. To test whether the communication between these two regions is necessary for the formation of the tone-shock association specifically in TFC, we chemogenetically silenced all SUB→RSP excitatory projections using designer receptors exclusively activated by designer drugs (DREADD) during training in both TFC and DFC.

We injected the DH of C57BL/6N male mice with Cre-independent inhibitory DREADD AAV8-hM4D_(Gi)_-mCherry virus and 6 weeks later microinfused the RSP with clozapine-N-oxide (CNO) through bilateral cannulas to locally block presynaptic release from RSP hM4D_(Gi)_-expressing axon terminals without modifying spiking activity of the cell soma (Figure 1A).^23^ Our chemogenetic manipulations targeted the SUB→RSP projections which are the major source of DH afferents to RSP (Figure S1A-B).^19^ In TFC trained mice, pre-training CNO infusions significantly impaired freezing during the tone and trace at tests, compared with vehicle control (Figure 1B), suggesting that SUB→RSP projections were necessary for the formation of tone-shock association. On the other hand, CNO injection did not affect freezing in DFC trained mice expressing Cre-independent inhibitory DREADD AAV8-hM4_(Gi)_-mCherry (Figure 1C). As expected, CNO infusions also had no effect on freezing in mice expressing the control virus AAV8-mCherry (Figure S2A). Additionally, at test, DFC trained mice did not freeze post tone, unlike mice trained in TFC paradigm, indicating that one-trial tone-shock pairing allows for clear differentiation between TFC and DFC at the behavioral level (Figure S2B). Together, these findings showed that SUB→RSP projections are necessary for TFC, but not DFC.

**Figure 1.**
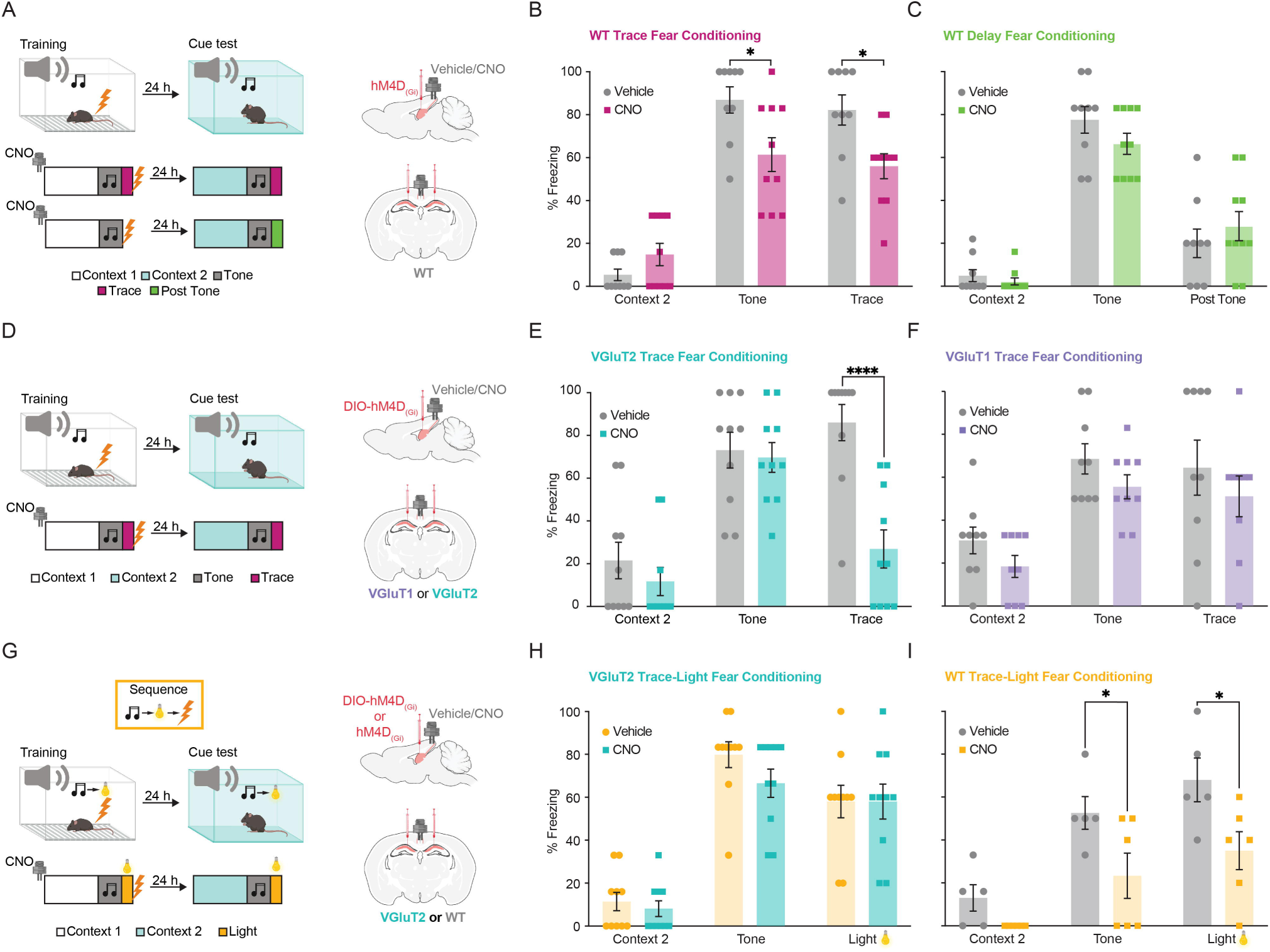
VGluT2^+^ SUB→RSP afferents are specifically required for processing a temporal trace separating a cue and a shock. **A**, **D**, **G** Experimental design of behavioral tasks. Diagrams depict virus infusion sites in DH and cannula placements for CNO injections in RSP, right. **B** When compared to vehicle, CNO injections 30 min before TFC training impaired freezing at test in response to both the tone and trace periods in mice receiving AAV-hM4D_(Gi)_ (n=9-10; two-way ANOVA with repeated measures; factor: treatment, P=0.0098, F_(1, 17)_= 8.444, factor: phase, P<0.0001, F_(2, 34)_= 67.41, factor: trial x phase, P=0.0084, F_(2, 34)_= 5.522). **C** CNO injection before training in Delay fear conditioning (DFC) did not affect freezing to tone or post-tone when compared to vehicle-injected mice (n=9-10; two-way ANOVA with repeated measures; factor: treatment, P=0.6738, F_(1, 17)_= 0.1835, factor: phase, P<0.0001, F_(2, 34)_= 97.84, factor: trial x phase, P=0.1764, F_(2, 34)_= 1.827). **E** In VGluT2-Cre mice CNO injection 30 min before training significantly impaired freezing during the trace but not tone compared to vehicle group (n=10; two-way ANOVA with repeated measures; factor: treatment, P=0.0132, F_(1, 18)_= 7.565, factor: phase, P<0.0001, F_(2, 36)_= 41.47, factor: trial x phase, P=0.0001, F_(2, 36)_= 12.00. **F** In VGluT1-Cre mice, terminal silencing with CNO did not affect freezing compared to vehicle controls (n=9; two-way ANOVA with repeated measures; factor: treatment, P=0.0931, F_(1, 16)_= 3.189, factor: phase, P<0.0001, F_(2, 32)_= 13.78, factor: trial x phase, P=0.9968, F_(2, 32)_= 0.003253) when compared to vehicle controls. **H** Injections of CNO (n = 10) before Trace-light conditioning (TLC) training did not affect freezing during either the tone or light periods in mice expressing inhibitory DREADD only in VGluT2^+^ SUB→RSP projections (n=10; two-way ANOVA with repeated measures; factor: treatment, P=0.2577, F_(1, 18)_= 1.366, factor: phase, P<0.0001, F_(2, 36)_= 52.27, factor: trial x phase, P=0.5714, F_(2, 36)_= 0.5685). **I** Injections of CNO before TLC training significantly impaired freezing to both tone and light when compared to vehicle in mice expressing inhibitory DREADD in all SUB→RSP projections compared to vehicle injected group (n=5-6; two-way ANOVA with repeated measures; factor: treatment, P=0.0087, F_(1, 9)_= 11.12, factor: phase, P<0.0001, F_(2, 18)_= 19.01, factor: trial x phase, P=0.3846, F_(2, 18)_= 1.008. Data presented as mean ± s.e.m. *P < 0.05, **P < 0.01, ***P < 0.001, ****P < 0.0001; NS, not significant; WT, wild type.

Our previous work demonstrated that the two excitatory SUB→RSP layer 3 projections differentially contribute to formation and persistence of recent and remote context memories.^19^ Thus, we next sought to determine, using selective chemogenetic silencing of VGluT1^+^ or VGluT2^+^ SUB→RSP terminals, whether this functional difference is extended to TFC. We injected the DH of male mice expressing Cre recombinase under control of either VGlut1 (VGluT1-Cre mice)^24^ or VGluT2 (VGlut2-Cre mice)^25^ promoter with Cre-dependent inhibitory DREADD AAV8-DIO-hM4D_(Gi)_-mCherry virus and 6 weeks later locally microinfused RSP with CNO through bilateral cannula before TFC training (Figure 1D). Silencing VGluT2^+^ terminals during training specifically impaired freezing during trace compared with control group (Figure 1E), while silencing VGluT1^+^ terminals during training had no effect on freezing (Figure 1F). Silencing SUB→RSP projections selectively during testing had the same effect (Figure S1C), indicating that exclusively activity of VGluT2^+^ terminals is necessary both for formation and recall of memory during trace. On the other hand, context memories (Figure S1D) required activity of VGluT1^+^ terminals, consistent with our previous findings.^19^ We excluded a sex effect by replicating these findings in female animals (Figure S1E-F).

To assess if the freezing impairment during trace caused by silencing VGluT2^+^ SUB→RSP terminals disrupted encoding of the temporal gap or event sequences, we used tone-light conditioning (TLC) paradigm. TLC is conceptually similar to TFC, except that the temporal gap separating two intermittent events in TFC is replaced by another neutral stimulus (flashing light) in TLC. Therefore, in TLC mice are presented with a sequence of three events (tone-light-shock) rather than two events separated in time as in TFC (tone-trace-shock). Following training, we quantified freezing in both TFC and TLC conditioned mice; first exposing them to a tone-trace, and then the following day to tone-light sequence. Both after TFC and TLC mice acquired fear conditioning to the tone, but freezing during the trace period was significantly higher in mice trained in TFC, whereas freezing during the light period was significantly higher in mice trained in TLC, suggesting that mice formed different associations (tone-trace-shock vs tone-light-shock) in these paradigms (Figure S2C). Next, we injected VGluT2-Cre mice with either Cre-independent or Cre-dependent inhibitory DREADD and 6 weeks later locally injected CNO in RSP to silence all excitatory or specifically VGluT2^+^ SUB→RSP afferents, respectively (Figure 1G). Inhibition of VGluT2^+^ SUB→RSP afferents did not affect freezing during TLC tests (Figure 1H), but inhibition all excitatory SUB→RSP terminals significantly impaired freezing during both tone and light presentation compared with controls (Figure 1I). This indicated that the trace freezing deficits induced by silencing VGluT2+ SUB→RSP projections were not a consequence of impaired processing of stimuli sequences, but likely reflected their inability to encode the temporal gap between tone and shock.

### Characterization of VGluT1^+^ and VGluT2^+^ neurons in the SUB**→**RSP circuit

Previous studies showed that throughout the brain VGluT1 and VGluT2 exhibit complementary expression,^26^ with neurons of the cerebellum, cortex and hippocampus primarily expressing VGluT1 positive, and subcortical brain areas mostly expressing VGluT2.^27^ To determine the neuronal origins of VGluT1 and VGluT2 projections, we used genetically modified VGluT1-Cre or VGlut2-Cre mice injected with Cre-dependent color flipping reporter retrograde virus to target these discrete populations. With this approach -where Cre^+^ population express GFP and Cre-population express TdTomato (Figure 2A)-we showed that both VGluT1 and VGluT2 neurons of the CA1 project to the SUB and that VGluT1 and VGluT2 neurons in the SUB project to the RSP (Figure 2B). Notably, when we injected Cre-dependent color flipping reporter virus with nuclear localization, we saw that CA1 and the SUB area have both VGluT1 and VGluT2 populations of neurons and that VGluT2 neurons are predominantly located in CA1 the deep layer, while in the SUB VGluT1 and VGluT2 neurons are spread heterogeneously (Figure 2C, D). Analysis of the online database Hippseq revealed that VGluT2 expression was limited to several SUB clusters (Fn1, Ly6g6e, S100b, Figure S5).^28,29^ Previous studies show that Fibronectin 1 (Fn1) is enriched in distal SUB, which provides main input to RSP,^30^ indicating that the VGlut2^+^ cells targeted in our study are likely Fn1 positive (Figure S5).

**Figure 2.**
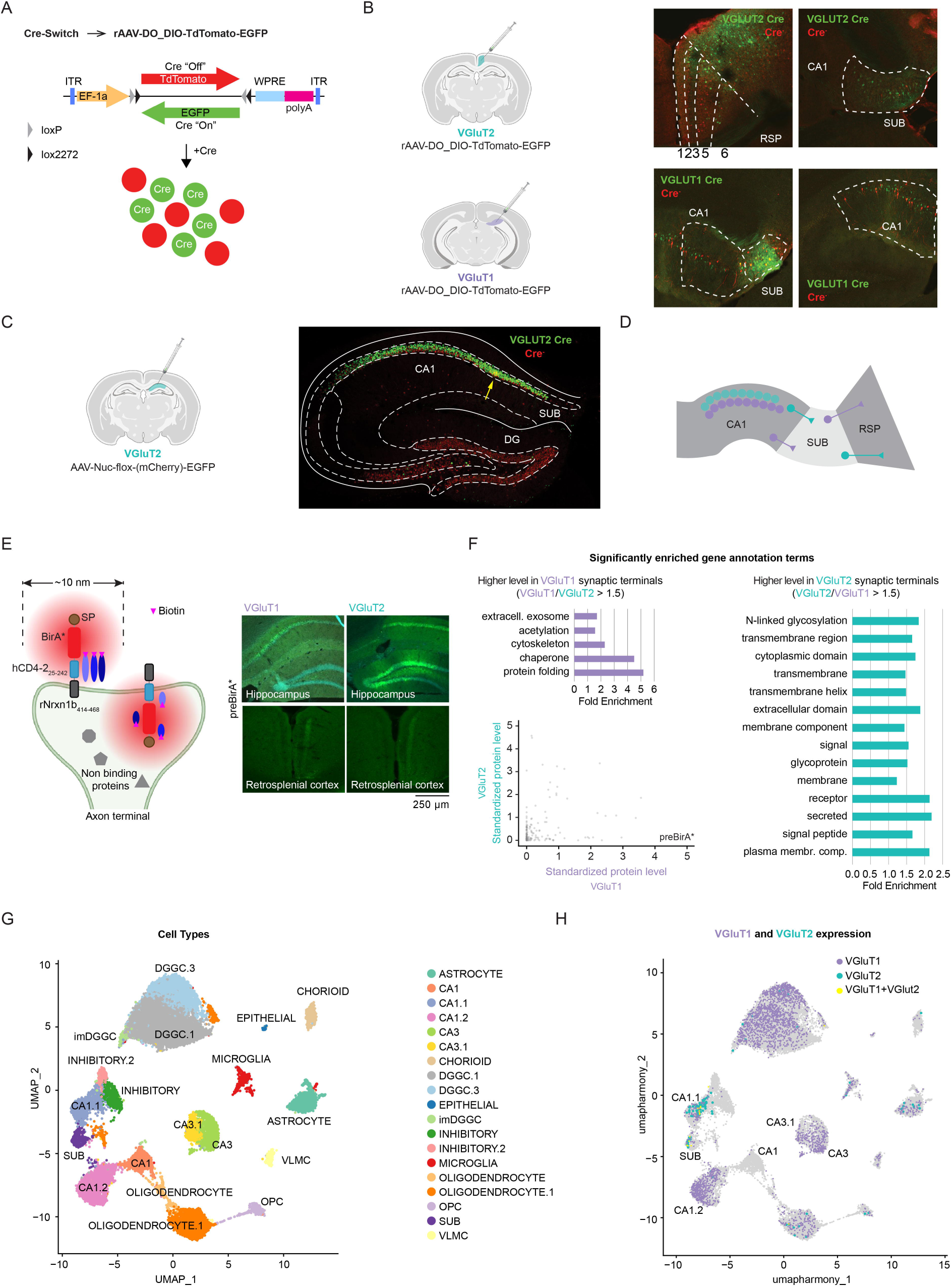
Characterization of VGluT1^+^ and VGluT2^+^ neurons in the SUB→RSP circuit. **A** Schematic of Cre-Switch viral vector (pAAV-Ef1a-DO_DIO-TdTomato_EGFP-WPRE). In Cre-negative cells, tdTomato is expressed, while in Cre-positive cells, Cre recombination inverts and excises the TdTomato cassette, leading to EGFP expression instead. **B** Top, retrograde Cre-dependent “color flipping” reporter (pAAV-Ef1a-DO_DIO-TdTomato_EGFP-WPRE) was injected into RSP of VGluT2-Cre mice. Red and green signals in the SUB show that both VGluT1^+^ and VGluT2^+^ population are projecting to the RSP. Bottom, retrograde Cre-dependent “color flipping” reporter was injected into SUB of VGluT1-Cre mice. Green and red signals in DH and SUB indicate presence of both VGluT1 and VGluT2 populations of neurons in both areas as well as presence of VGluT1^+^ and VGluT2^+^ CA1→SUB projections. **C** Cre^+^ (VGluT2^+^, green) and Cre-(red, presumably VGluT1^+^) hippocampal neurons visualized after injection of the Cre-dependent color flipping nuclear reporter (pAAV8-EF1a-Nuc-flox(mCherry)-EGFP) into the DH. **D** Summary diagram of VGlut1 and VGluT2 expression based on results from the Cre-switch reporter injections. **E** Left, schematic of proximity biotinylation assay. Right, injection sites for preBirA* in DH and corresponding GFP signal in DH and RSP. **F** Right, cytoBirA* was injected into DH of naïve VGluT1-Cre and VGluT2-Cre mice, biotinylated proteins were pulled down, and quantified from RSP. Left and up, KEGG terms of differentially expressed proteins in VGluT1^+^ or VGluT2^+^ biotinylated terminals. Bottom left, After injection of the virus expressing BirA*-Pre, the levels of biotinylated proteins from VGluT1 RSC were dissimilar, as indicated by lack of correlation, suggesting differences in their presynaptic proteomes. **G** Main cell populations identified using known neuronal and non-neuronal cell markers. **H** VGluT1 and VGluT2 expression across cell types reveals largely non-overlapping populations.

To examine the proteins expressed in the SUB→RSP terminals, we used in vivo proximity biotinylation in combination with mass spectrometry (MS)-based proteomics analysis^31,32^ (Figure S3A; see Methods). Similar to the tracing approaches, EGFP co-expressed by preBirA* AAV constructs localized in RSP layer 3 (Figure 2E). To investigate the proteomic diversity of VGluT1 and VGluT2 presynaptic terminals, we employed tandem mass tag (TMT)-based quantitative mass spectrometry (MS) to directly compare their proteomes. Specifically, VGlut1-Cre and VGlut2-Cre mice were injected with either AAV-FLEx-preBirA* or AAV-FLEx-cytoBirA*. Proteins exhibiting a preBirA* to cytoBirA* ratio grater then 1.5 were considered enriched and thus classified as presynaptic terminal proteins. We mined the VGluT1^+^ and VGluT2^+^ SUB→RSP terminal proteomic datasets using the online Database for Annotation, Visualization, and Integrated Discovery (DAVID). As indicated by KEGG analysis, the proteomes of VGluT1^+^ and VGluT2^+^ RSP terminals differed from one another under baseline conditions (Figure S3B). VGluT2^+^ terminal proteome showed significantly enriched KEGG terms involved in neurotransmitter release and synaptic plasticity (Endocytosis, SNARE interactions in vesicular transport, Ribosome, Long-term potentiation and Axon guidance), and various neuromodulatory signaling and co-transmitter phenotype (Endocannabinoid, GABAergic, Dopaminergic and Cholinergic synapse),^26^ activity-dependent modulation and involvement in learning and memory processes (Figure S3B). Overall, differentially expressed proteins in VGluT2 biotinylated terminals were glycoproteins, membrane, receptor, and transmembrane proteins involved in signaling (preBirA*/cytoBirA* > 1.5 fold), while the greatest changes (preBirA*/cytoBirA* > 1.5 fold) in VGluT1 terminals were in proteins involved in exosome function, acetylation, cytoskeletal dynamics, chaperone function, and protein folding (Figure 2F). We correlated the levels of biotinylated proteins pulled down from VGluT1^+^ and VGluT2^+^ terminals to control for specificity of the proteomic differences. Samples from mice injected with preBirA* (Figure 2F, down left) were highly divergent (R^2^ = 0.03), reflecting substantial differences in their presynaptic terminal proteomes. In contrast, in mice injected with the control virus cytoBirA* (Figure S3C), protein abundances showed a strong positive correlation (R^2^ = 0.82), indicating similar protein profiles due to non-specific biotinylation of cytoplasmatic proteins. Together these data shows that VGluT1^+^ and VGluT2^+^ SUB→RSP have distinct presynaptic proteomic signatures, indicating a specialized role in synaptic function. VGlut2^+^ proteome suggest a more modulatory and dynamic role, while the VGluT1^+^ terminals show profile associated with structural support and protein synthesis.

To further explore VGluT1^+^ and VGluT2^+^ neuronal and non-neuronal hippocampal populations, we used our previously published dataset for which we dissected the DH and used 10×Genomics platform to perform single-nucleus RNA-sequencing (snRNA-seq).^33^ An unsupervised algorithm run on the snRNA-seq dataset identified 30 clusters (Figure S4A). Using canonical markers and existing hippocampal databases (see Methods), we identified CA1 neurons (3 clusters), SUB neurons (1 cluster) CA3 neurons (2 clusters), dentate gyrus granule cells (DGGC) (4 clusters), interneurons (2 clusters), and different non-neuronal cells (8 clusters; Figure 2G and S4A). snRNA-seq data showed that VGluT1 and VGluT2 were present in non-overlapping cellular populations (Figure 2H and S4B), which concurs with our tracing experiments. A small subset of cells in the CA1.1 and SUB clusters was both VGluT1 and VGluT2 positive, a population also seen in DH of mice injected with Cre-dependent color flipping nuclear reporter (Figure 2C, right, indicated with yellow arrow). Overall, we demonstrate that VGluT1 phenotype is most prevalent in the majority of hippocampus, with exception of CA1 and SUB, which contain mostly non-overlapping VGluT1^+^ and VGluT2^+^ populations. Both CA1 and SUB outputs are VGluT1^+^ and VGluT2^+^, however, in the CA1 region VGluT1^+^ and VGluT2^+^ populations are organized into deep and superficial layers, while the SUB shows a more heterogenous distribution of VGluT1^+^ and VGluT2^+^ populations (Figure 2D).

### VGluT2^+^ SUB→RSP afferents have distinct response dynamics dependent on TFC phase

To assess activity of VGluT2^+^ SUB→RSP afferents during TFC, we used fiber photometry to measure the populations’ activity through changes in Ca^2+^ activity. We injected mice with Cre-dependent calcium indicator AAV-DIO-GCaMP7s in DH of VGluT2-Cre mice, implanted with a fiber targeting the SUB→RSP layer 3 projections and recorded the changes in the fluorescence signal -as an indicator of bulk calcium activity-across 3 trials of training and testing days (Figure 3A). Consistent with previous studies we found that every presentation of a foot shock during training elicited a large, sustained increase in the GCaMP fluorescence (Figure 3B, D).^9^ We compared average z score for signals obtained from the RSP terminals originating in the SUB across different phases of training and found that fluorescent signal increased during first 15 s of the tone followed by a dip during trace with repeated tone-shock pairing (Figure 3 E). Area under the curve (AUC) showed a similar pattern of signal dynamics, where onset of tone caused a significant increase in the GCaMP signal at the second and third presentation of tone, that was followed by the negative-going responses (“dips”) during trace. Dip in GCaMP signal during trace was significantly amplified at third tone presentation (Figure S6A). During the cue test, average z scores showed differences in bulk calcium activity during the first 15 s of the tone and 15 s of trace at first and last trial (Figure 3C, E). On the testing day, a significant increase of the GCaMP signal at the tone onset was elicited at the first presentation to tone, followed by a moderate negative-going response increase (Figure 3C, E). While the signal increase at the tone onset was only present at the first tone presentation, dip during trace period reemerged at third tone presentation (Figure 3C, E). AUC comparison showed similar signal dynamics (Figure S6B). Together these results show that VGluT2^+^ SUB→RSP projection response dynamic is characterized by overshoot at tone onset followed by the signal dip during trace formed during memory acquisition and replayed at first memory recall.

**Figure 3.**
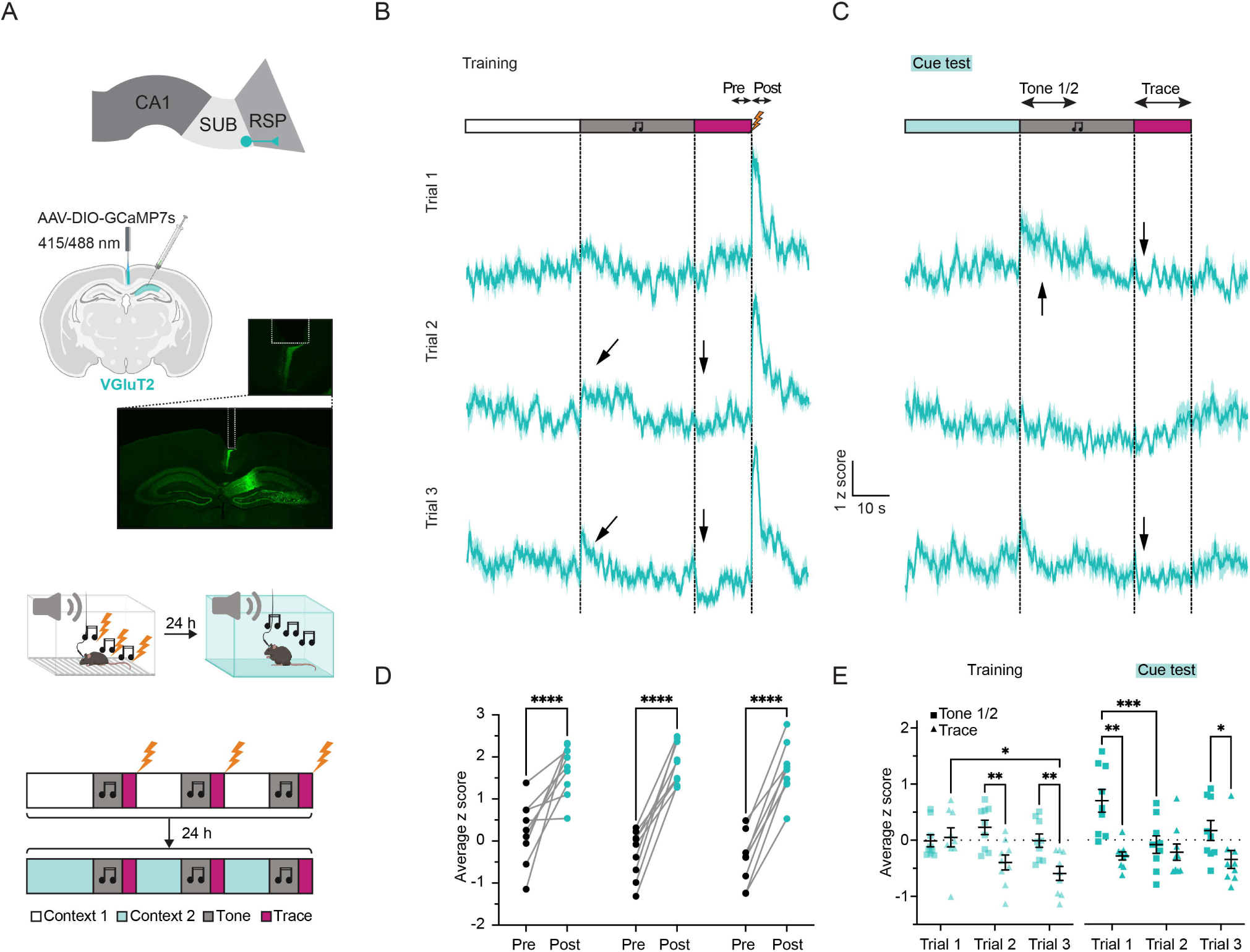
VGluT2^+^ SUB→RSP afferents have distinct response dynamics dependent on TFC phase. **A** Top, representative image of viral expression and fiber track, together with the diagram indicating position of injection and fiber placement. Bottom, Experimental design for fiber photometry recordings, bottom. **B C** Average GCaMP fluorescence traces during 3 CS-US presentations on training day (**B**) and during 3 CS presentations on test day (**C**) recorded from RSP VGluT2^+^ terminals of projections originating in SUB. Arrows indicate significant increases and decreases of signal. **D** Increase in average z score following shock exposure (n=9; two-way ANOVA with repeated measures; factor: trial, P=0.2195, F_(2,32)_= 1.591, factor: phase, P<0.0001, F_(1, 16)_= 97.98, factor: trial x phase, P=0.1259, F_(2, 32)_= 2.212). **E** Average z-score at tone onset and during trace on training (left) and testing (right) day. Training Tone vs Trace (n=9; two-way ANOVA with repeated measures; factor: trial, P=0.0689, F_(1.927, 30.84)_= 2.951, factor: phase, P=0.0021, F_(1, 16)_= 13.40, factor: trial x phase, P=0.0258, F_(2, 32)_= 4.107). Cue test Tone vs Trace (n=9; two-way ANOVA with repeated measures; factor: trial, P=0.0355, F_(1.910, 30.56)_= 3.795, factor: phase, P=0.0029, F_(1, 16)_= 12.37, factor: trial x phase, P=0.0165, F_(2, 32)_= 4.679). Data presented as mean ± s.e.m. *P < 0.05, **P < 0.01, ***P < 0.001, ****P < 0.0001; NS, not significant.

### Changes in response dynamics of VGluT2^+^ SUB→RSP afferents are specific to learning of tone-shock association

To exclude the possibility that the observed response dynamics is a non-associative effect of tone-induced arousal rather than consequence of associative learning, we recorded GCaMP signals from VGluT2^+^ SUB→RSP afferents in mice undergoing pseudoconditioning.^34^ TFC and pseudoconditioning differ by a key feature - timing between shock and tone; in pseudoconditioning shock and tone are presented temporally unpaired, preventing the formation of a predictive relationship between two stimuli. Thus, if the response dynamic is a consequence of the learned tone-trace-shock association, it will not appear at tone presentation during pseudo fear conditioning. At training, all three shock presentations elicited a response, indicating that shock response during training is not specific for tone-shock association (Figure 4A-C). Tone presentation both during training and testing did not cause changes in GCaMP fluorescent signal suggesting that the pattern of activity observed during TFC is a consequence of tone-trace-shock association (Figure 4D, E). Taken together, these data suggest that differences in activity dynamic of VGluT2^+^ SUB→RSP afferents encode association of the tone and shock that are separated by a time gap.

**Figure 4.**
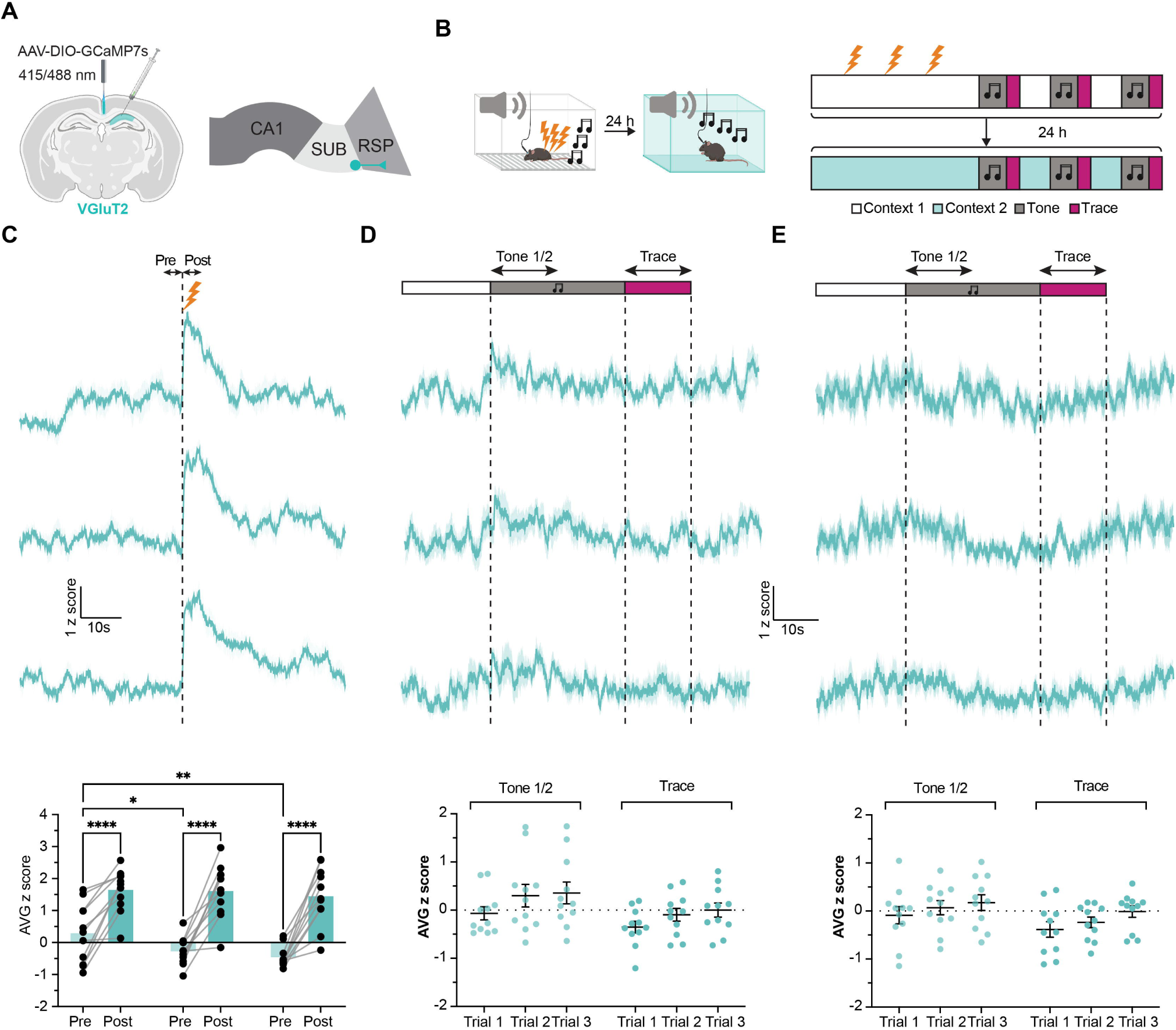
Changes in response dynamics of VGluT2^+^ SUB→RSP afferents are specific to learning of tone-shock association. **A** Experimental diagram detailing GCaMP7s AAV injection in the DH and fiber placement in the RSP, enabling recording of calcium signals from SUB→RSP terminals. **B** Schematic of the unpaired fear conditioning paradigm. **C** Top, averaged GCaMP fluorescence traces during 3 unpaired shock presentations on training day. Bottom, significant differences were found in average z score between pre shock and post shock period (n=11; two-way ANOVA with repeated measures; factor: trial, P=0.0199, F_(2, 40)_= 4.324, factor: phase, P<0.0001, F_(1, 20)_= 48.23, factor: trial x phase, P=0.1750, F_(2, 40)_= 1.8210) **D** Top, averaged GCaMP fluorescence traces during 3 unpaired tone presentations on training day. Bottom, no significant differences were found in average z score between first half of the tone and trace (n=11; two-way ANOVA with repeated measures; factor: trial, P=0.0988, F_(2, 40)_= 2.454, factor: phase, P=0.0068, F_(1, 20)_= 9.090, factor: trial x phase, P=0.9558, F_(2, 40)_= 0.0453) **E** Top, averaged GCaMP fluorescence traces during 3 unpaired tone presentations during the cue test. Bottom, no significant differences were found in average z score between first half of the tone and trace (n=11; two-way ANOVA with repeated measures; factor: trial, P=0.1915, F_(2, 40)_= 1.723, factor: phase, P=0.0018, F_(1, 20)_= 12.92, factor: trial x phase, P=0.9272, F_(2, 40)_= 0.0758)

### CA2 and CA3 responses during TFC differed from the responses of the SUB**→**RSP projections

Post-shock increase in GCaMP signal has been previously observed in CA1.^9^ Here, we showed that VGluT2^+^ SUB→RSP projections during training show a large, sustained increase in the GCaMP fluorescence in response to foot shock (Figure 3B, D) indicating input signal from the CA1. Considering that we observed signal dynamic at the VGluT2^+^ SUB→RSP afferents that was different from previously reported CA1 dynamic, we sought to test if CA2 and CA3, which are major CA1 inputs, additionally project to the SUB and if added complexity of the VGluT2^+^ SUB→RSP projection signal dynamics might correlate with CA2 or CA3 activity. Using Grik4-Cre and Amigo-Cre mice that specifically limits viral expression to CA3 hippocampal subarea and pAAV-hSyn-FLEx-mGFP-2A-Synaptophysin-mRuby viral vector we show that in addition to CA1, the CA2 and CA3 also send projections to SUB (Figure S8-9). Both CA2 and CA3 showed increase of GCaMP signal following shock during training (Figure 5C) but no significant responses to the tone both during training and testing (Figure S7). While both areas responded to shock, CA3 response diminished with repetitive trials, unlike CA2 (Figure 5D). During training and testing, CA2 did not respond to the tone presentations (Figure S7A, B). On the other hand, at recall GCaMP signal in the CA3 region significantly increased during trace following the first presentation of the conditioned stimulus (tone) (Figure 5F and Figure S7C, D). Taken together, these data indicate apart from their shock responses, the activity of the CA2 and CA3 neurons did not resemble the patterns observed in the VGluT2^+^ SUB→RSP projections.

**Figure 5.**
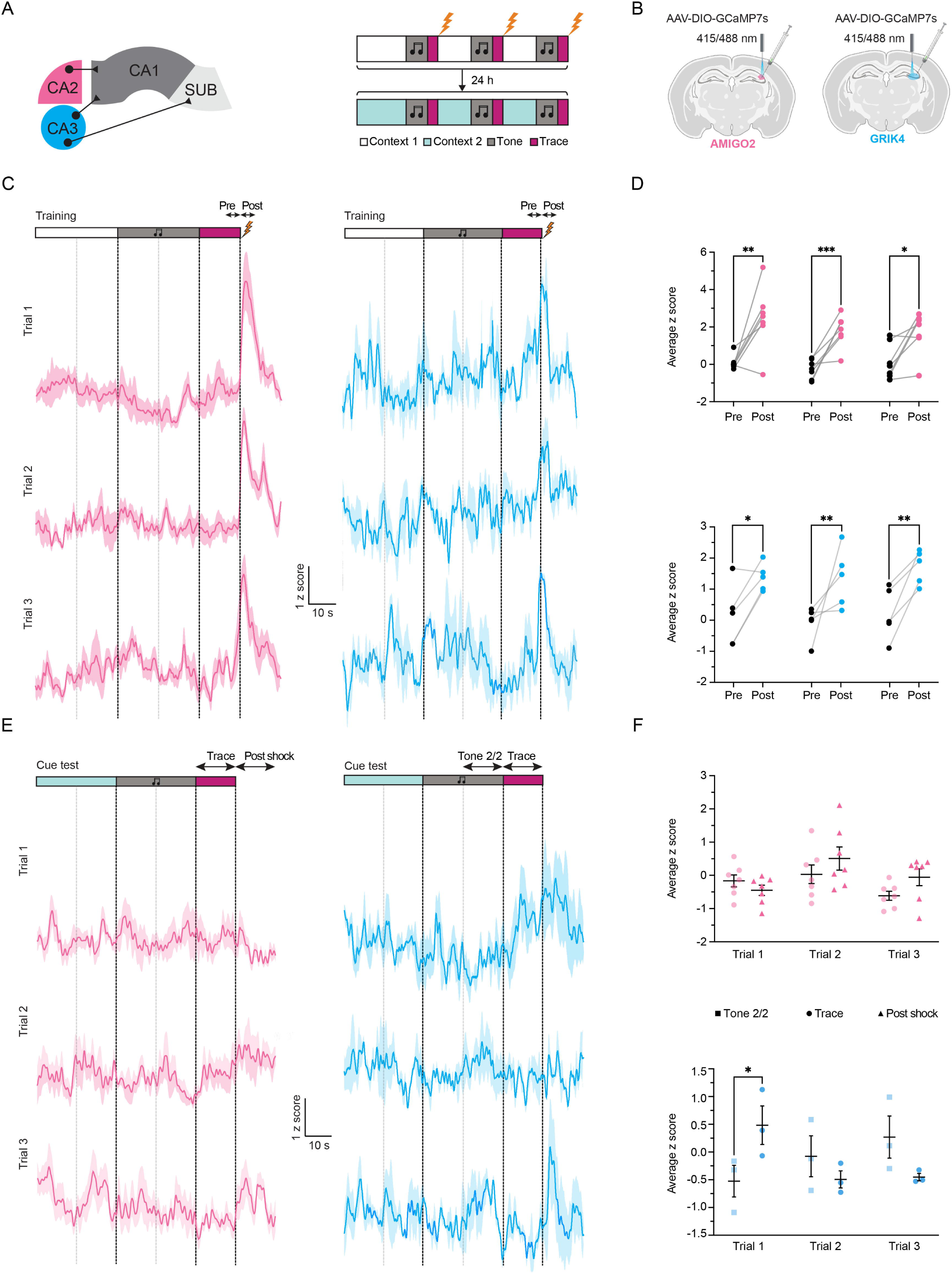
CA2 and CA3 responses during TFC differed from the responses of the SUB→RSP projections. **A** Left, diagram showing the connectivity of hippocampal subfields CA2 and CA3. Right, experimental design for fiber photometry recordings. **B** Schematics indicating position of injection and fiber placement in Grik4-Cre and Amigo2-Cre mice, targeting CA3 and CA2 areas of the DH, respectively. **C** Average GCaMP fluorescence traces during 3 CS-US presentations on training day recorded from Amigo2 positive CA2 cell bodies and fibers (left) and CA3 cell bodies (right). **D** Top, repeated significant increase in average z score following shock exposure at CA2 (n=7; two-way ANOVA with repeated measures; factor: trial, P=0.1835, F_(1.693, 20.31)_= 1.869, factor: phase, P=0.0007, F_(1, 12)_= 20.53, factor: trial x phase, P=0.3200, F_(2, 24)_= 1.195). Bottom, similar increase was also observed in the CA3 (n=5; two-way ANOVA with repeated measures; factor: trial, P=0.6647, F_(2, 16)_= 0.4190, factor: phase, P=0.0006, F_(1, 8)_= 29.14, factor: trial x phase, P=0.9169, F_(2, 16)_= 0.0873) **E** Average GCaMP fluorescence traces during 3 CS-US presentations on test day recorded from CA2 cell bodies and fibers (left) and CA3 cell bodies (right). **F** Top, no significant changes were found in average GCaMP z score at trace and post shock in CA2 (n=7; two-way ANOVA with repeated measures; factor: trial, P=0.0223, F_(2, 24)_= 4.476, factor: phase, P=0.2533, F_(1, 12)_= 1.440, factor: trial x phase, P=0.1475, F_(2, 24)_= 2.075). Bottom, in the CA3, a significant trail x phase interaction was found (Tone 2/2 vs Trace, n=3; two-way ANOVA with repeated measures; factor: trial, P=0.6028, F_(2, 8)_= 0.5396, factor: phase, P=0.8854, F_(1, 4)_=0.0236, factor: trial x phase, P=0.0240, F_(2, 8)_= 6.158). Data presented as mean ± s.e.m. *P < 0.05, **P < 0.01, ***P < 0.001, ****P < 0.0001; NS, not significant.

## Discussion

We demonstrated that VGluT2^+^ SUB→RSP projections significantly contribute to the associative and temporal components of temporal associative memories, which were presented to the RSP as an integrated pattern of activity acquired during training and reinstated at retrieval. This was indicated by the observations that concurrent inhibition of VGluT2^+^ and VGlutT1^+^ SUB→RSP projections impaired tone-shock associations whereas inhibition of the VGluT2^+^ SUB→RSP projections selectively impaired freezing during the trace interval. During TFC, VGluT2^+^ SUB→RSP projections acquired pattern of bulk calcium activity involving a transient increase of the calcium signal at tone onset, followed by suppression during the trace interval. Such activity patterns were not seen in the CA2 and CA3, as shown here, and in CA1, as reported previously,^9^ suggesting that integration of associative and temporal components of TFC occurred at the SUB or even at the level of the synaptic terminals (Figure 6).

**Figure 6.**
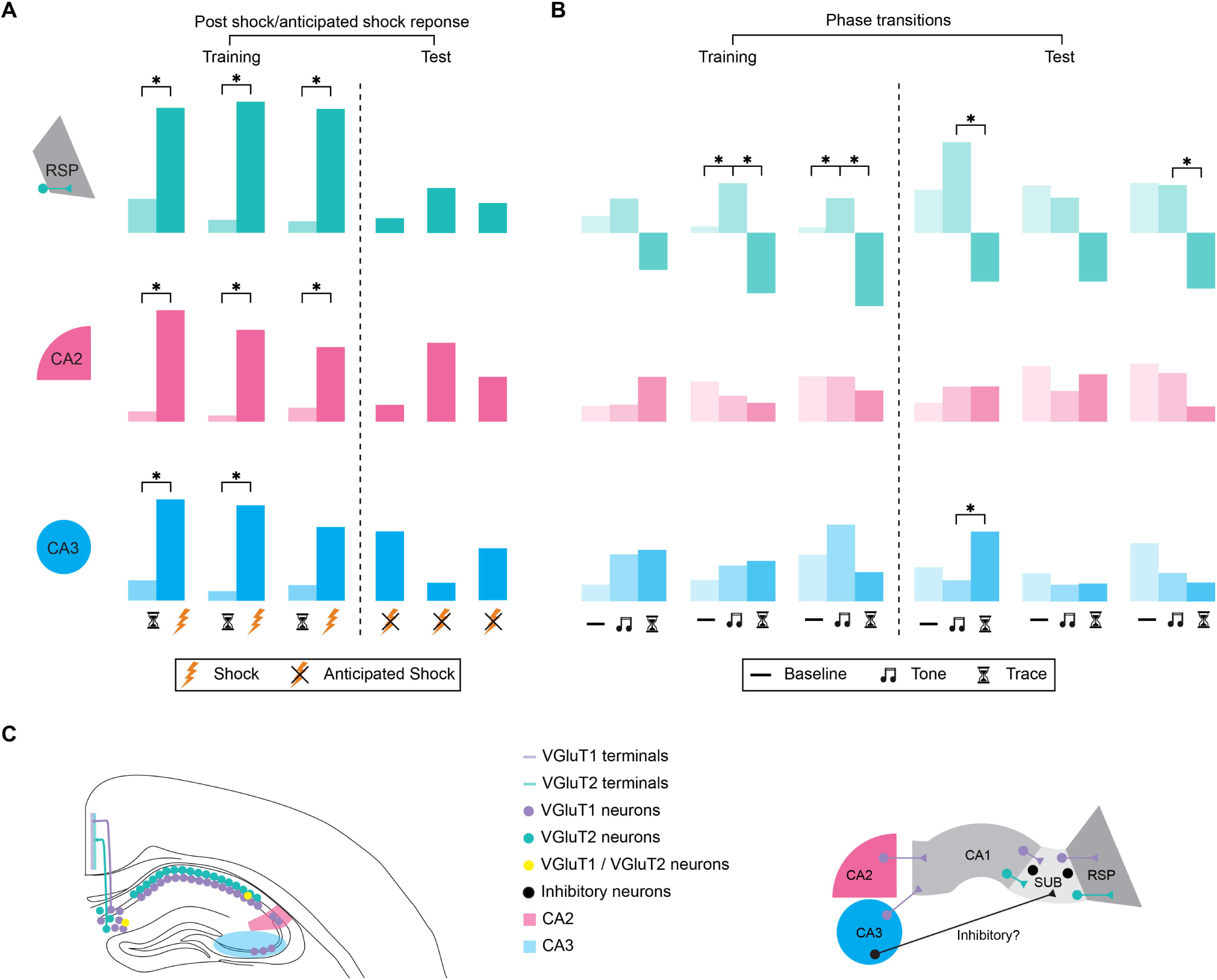
Summary of hippocampal circuit activity dynamic underlying TFC. **A** Schematics of the key findings from fiber photometry recordings of VGluT2^+^ SUB→RSP projections and hippocampal subregions CA2 and CA3 during training (just before and at shock occurrence) and testing (in anticipation of shock that occurred at training but is omitted at testing). All three areas recorded show activation in response to shock during training. At testing, “anticipated shock” refers to the time point corresponding to shock delivery during training session but where no shock was administered. Each region shows distinct dynamics related to this expected but omitted aversive stimulus. **B** Schematic depicting learning-dependent changes in activity in VGluT2 SUB→RSP terminals and CA2/CA3 subregions during training and memory retrieval. During training, RSP terminals show increased activation at tone onset followed by a signal “dip” during the trace interval. At memory retrieval, this biphasic response in RSP terminals is observed only during the first CS presentation, while CA3 displays increased activity during the trace period—reflecting anticipatory processing. Asterisks indicate significant effects. **C** Left, schematic highlighting VGluT1and VGluT2 expression pattern in DH, informed by our snRNA seq data and previously published literature. Right, simplified model of dorsal hippocampal connectivity based on our viral tracing experiments and previously published data.

We previously demonstrated that the formation of associative contextual memories predominantly depends on the VGluT1^+^ SUB→RSP projections while VGluT2^+^ projections contributed to memory persistence.^19^ Here we found a similar functional divergence but with respect to the encoding of the temporal trace, which solely depended on the activity of VGluT2^+^ SUB→RSP projections. This was indicated by the effects of chemogenetic inhibition of these projections, which specifically impaired freezing during the trace period, but not during the tone, or sequences of tone-light-shock pairings lacking a temporal gap.

Early postnatal excitatory circuits rely on VGluT2 activity, but, as hippocampal, cortical, and cerebellar circuits mature, neurons switch from VGluT2 to VGluT1 phenotype.^35^ The CA1, the SUB, and a couple of other brain areas are unique in that their neurons express both vesicular transporters. While several findings suggested that VGluT1 and VGluT2 confer different functional properties to excitatory projections, such as different synaptic vesicle dynamics and glutamate release probability, ^36,37^ it has remained unclear whether they stem from discrete or overlapping neuronal populations. Here we showed, using both viral and single-nucleus RNA sequencing approaches, that only a small subset of CA1 and SUB neurons contains both transporters, whereas most are positive for either VGluT1 or VGluT2. In the CA1, we show that VGluT2^+^ neurons were predominantly located in the developmentally older,^38^ deep layer, suggesting that these neurons, rather than switching to VGluT1, retained their early developmental VGluT2 phenotype. The segregation of VGluT1 and VGluT2 across deep and superficial layers could contribute to their robust differences in firing rates and burst frequency,^39^ as well as in their unique contributions to spatial and temporal associative memories. The VGluT neuronal phenotypes in the distal SUB, the primary source of RSP projections, appeared less segregated. Based on our analysis of publicly available databases showing co-expression of VGluT2 and fibronectin 1 (Fn1)^28^, it is likely that our manipulations primarily targeted VGluT2^+^ Fn1^+^ neurons of the distal SUB. In addition to their different cellular origins, excitatory SUB→RSP projections also showed significant differences in their synaptic proteomes, with VGluT2^+^ presynaptic terminals showing enrichment in neuromodulatory and SNARE-mediated vesicular transport proteomes relative to VGluT1^+^ synapses. The overall greater proteomic variability of VGluT2^+^ terminals may be particularly adept at modulating cortical networks in response to dynamic sensory inputs (e.g., TFC versus contextual fear conditioning), thereby fine-tuning cortical circuits for specific behavioral outcomes.

It is currently believed that temporal associations are encoded through three interacting processes: trace holding (recent sensory input), temporal expectation (prediction of forthcoming stimuli), and explicit timing (estimation of stimulus onset relative to prior cues).^40^ According to prevailing models, sustained neural representations of the conditioned stimulus (tone in the case of TFC) across the trace interval are required for the association of temporally discontiguous events.^41^ Single cell analyses of the activity of DH and SUB neurons have not provided conclusive evidence on learning-related activity patterns coding for the different stimuli and the temporal gap between them. Deep-brain calcium imaging identified a subgroup of neurons both in the CA1 and the SUB, that maintained tone-evoked activity during the trace period at memory acquisition and consolidated this plasticity after learning in CA1 (but not SUB), becoming increasingly active in anticipation of shock during the tone-trace interval, an activity correlated with later fear retrieval. Based on these findings, it was suggested that the SUB temporarily supports stimulus maintenance, while CA1 stably encodes this information for long-term memory formation.^13^ Importantly, the study used AAV (serotype 9) driven by the CaMKIIα promoter for expression of the calcium sensor which is known to target both excitatory pyramidal neurons and subsets of GABAergic interneurons,^42^ suggesting that some of the calcium signals could potentially reflect activity of local SUB interneurons. Other studies showed more nuanced responses of CA1 neurons. Two-photon imaging has identified a sparse group of cue-activated CA1 neurons displaying stochastic dynamics across trials, suggesting that trace is not bridged by persistent activity but rather widespread shifts in neuronal ensemble activity,^43^ whereas single unit recordings showed increased firing only at the first presentation of the tone.^44^

Our analysis of bulk synaptic activity, revealed by calcium responses in VGluT2^+^ SUB→RSP synapses, revealed a different activity pattern: an increase in fluorescent signal at tone onset, followed by a pronounced signal dip during the trace interval. These responses were acquired during training, reconstructed during the first memory test, and not found after pseudo conditioning, demonstrating their specificity for temporal associative learning as well as temporal coding. Current views assume that the association between two events separated in time is simply strengthened by sustained excitatory firing during trace, with inhibition serving merely as a dampening force.^40,41^ Our findings indicate that, rather than simply reflecting a drop in excitatory drive, this dip may instead mark a temporally structured inhibition that regulates how and when subicular output modulates RSP activity required for temporal coding.^17^ Feedforward inhibition of SUB VGluT2^+^ neurons is a likely mechanism for the decrease of their activity during the temporal interval within which associative learning can take place, however, it is unclear at this time whether such inhibition is provided by local microcircuits or from long-range inhibitory projections marked by neuropeptide Y (NPY), somatostatin (SOM), and muscarinic acetylcholine receptor type 2 (M2R).^45^ These pathways could temporally inhibit SUB output, at least during the trace memory test.

Interestingly, recordings of calcium signals from individual DH subfields that innervate the CA1 and SUB, apart from shock responses, did not show cue- or trace-specific patterns during TFC as found in the VGluT2^+^ SUB→RSP projections. Our findings with CA2 and CA3 are consistent with the CA1 model proposing that TFC induces latent memory traces that are stabilized post-shock through sharp-wave ripples,^9^ which are provided by CA2 and CA3.^46–48^ Nevertheless, none of the activity patterns seen in the VGluT2^+^ SUB→RSP terminals was observed in the individual hippocampal subfields. There are several possible explanations for the differences in the activity patterns detected in SUB inputs and outputs. It is possible that synaptic population activity reveals TFC-specific patterns better than neuronal population activity. The predictive activity pattern at VGluT2^+^ SUB→RSP terminals implied the presence of circuit-level computation within the SUB. This observation aligns with prior findings that designate the SUB as a critical hub for transforming and routing hippocampal output.^49^ Indeed, large scale optogenetically targeted electrophysiological recordings show that dorsal SUB neurons conveyed comparable or greater information per unit of time compared to CA1 neurons, and depending on information type, SUB conveyed information out either uniformly or selectively to specific projection targets, with their firing precisely controlled by theta oscillations and SWR.^49^ Such mechanism could enable flexible modulation of downstream cortical networks in a manner consistent with dynamic memory demands. Alternatively, TFC-specific activity patterns in VGluT2^+^ SUB→RSP terminals may not be directly correlated with SUB neuronal activity,^13^ but instead generated through plasticity at discrete SUB→RSP synapses.

There is emerging support for the view that separate mechanisms contribute to the processing of different components of human episodic memories, with different developmental trajectories for memories of facts, space, and time.^50,51^ The functional differentiation of circuits identified in our work might thus be applicable to human memory processes from early development throughout adulthood. Moreover, our findings suggest that temporal coding in TFC may emerge not from excitatory persistence but from a circuit-level interplay of excitation and well-timed inhibition, coordinated along the hippocampus–SUB–RSP axis. Future studies are required to delineate the sources of inhibitory inputs to the SUB and to define how its output signals are integrated within downstream target areas.

### Limitations

The current study recorded bulk calcium signal activity because our main interest was in the synaptic activity patterns conveyed from SUB to RSC. However, since we recorded from the CA2 and CA3 cell bodies we recorded activity of all neurons in the CA2 and CA3 area. Such approach could mask the detection of activity of individual neurons that projects to SUB, which may exhibit finer, more specific patterns of activity, and could overlook transient or subtle dynamics that could provide additional insight into the underlying processes of temporal memory consolidation.

## Resource availability

Lead contact: Requests for further information and resources should be directed to and will be fulfilled by the lead contact, Ana Cicvaric.

## Materials availability

This study did not generate new unique reagents.

## Data and code availability

All results of synaptic proteomes analysis have been deposited through an interactive portal (https://proteome-phenome-atlas.com/) and are publicly available as the date of publication. To generate snRNA seq graphs this paper analyzes existing, publicly available data, accessible at NCBI Gene Expression Omnibus database under accession GSE254780 and source code available at https://github.com/RadulovicLab/Nature-2024. All other data reported in this paper and any additional information required to reanalyze the data reported in this paper is available from the lead contact upon request. All other software and analytical methods used in this study are publicly available, as listed in the key resources table.

## Acknowledgements

This work was funded by NIMH grants MH108837 and MH078064 to J.R., J 4271 FWF (Austrian Science Fund) to A.C., NARSAD Young Investigator Grant to YZ.W., Individual Biomedical Research Award from The Hartwell Foundation to J.N.S. We thank Genetics and Computational Genomics Cores at Albert Einstein College of Medicine, especially Junya Zhang, for their help in data analysis. Illustrations of nonscientific data were created with BioRender.com through an institutional license.

## Author contributions

JR and AC designed the study. AC, TEB and JR wrote the original draft. YZW, VJ, NK and JNS performed proteomic experiments. EMW performed snRNA seq experiments. AC and TEB performed fibephotometry experiments. NY, AC and TEB performed tracing experiments. Lr, VG, JR and AC performed behavior tests and data analysis. HZ, ZP and VP. YZW, JNS, HZ, EMW and ZP assisted in manuscript revision. JR and AC obtained funding.

## Supplemental information

Figures S1-S9

Supplementary Tables 1-2.

## STAR Methods

### Animals

We used male and female C57BL/6N mice, vGlut1-Cre^24^, vGlut2-Cre^25^, Amigo-Cre^52^ and Grik4-Cre^53^ mice, as described in detail recently.^54^ Wild type C57BL6/J mice were purchased from Harlan, Indianapolis, IN. All Cre mouse lines were obtained from the Jackson Laboratory (Bar Harbor, ME). The VGluT1-Cre mouse line, also known as Slc17a7-IRES2-Cre or Vglut1-IRES2-Cre-D, was created by the Hongkui Zeng lab, Allen Institute for Brain Science^24^, the VGluT2-Cre knockin mice, also known as Slc17a6tm2(cre) and Lowl or VGlut2-ires-Cre, was generated as described previously^25^. The Amigo2-Cre mouse line, also known as B6.Cg-Tg(Amigo2-cre)1Sieg/J, was created in the laboratory of Dr. Steven A. Siegelbaum, as described previously.^52^ We used this mouse line to restrict the viral expression to CA2 hippocampal subregion as in adult Amigo2-Cre mice the *cre* is expressed predominantly in the pyramidal neurons of the CA2. For fiberphotometric recordings from the CA3 region we used Grik4-Cre, also known as C57BL/6-Tg(Grik4-cre)G32-4Stl/J, that have *cre* expression is restricted to pyramidal neurons of the CA3, created in the laboratory of Dr. Susumu Tonegawa, as described previously.^53^

Mice were bred, genotyped (using primers reported on the Jackson Laboratory website) and used for experiments at the age of 8 weeks. Typically, we obtained 4-6 litters/breeding cycle with 5-8 mice/litter with similar distribution of males and females. All litters were used for behavioral experiments and randomly allocated mice were used for tracing, behavioral and proteomic studies. The mice were maintained under standard housing conditions (12 h/12 h light dark cycle with lights on at 7 a.m., temperature 20-22 °C, humidity 30-60%) in our satellite behavioral facility. All animal procedures used in this study were approved by the Northwestern University’s Animal Care and Use Committee (protocols IS00002463 and IS00003359) and Albert Einstein’s Animal Care and Use Committee (protocols 00001289 and 00001268) in compliance with US National Institutes of Health standards.

### Stereotaxic surgeries and infusions of viral vectors and drugs

Mice were anesthetized with 1.2% tribromoethanol (vol/vol, Avertin) for viral vector intracranial infusion and cannula implantation. The Cre-dependent color-flipping retrograde reporter pAAV-Ef1a-DO_DIO-TdTomato_EGFP-WPRE-pA^55^ (gift from Bernardo Sabatini, Addgene plasmid Cat# 37120) was injected unilaterally in RSP or SUB (RSP: 1.80 mm posterior, ±0.40 mm lateral, 1.00 mm ventral to bregma and SUB: 3.52 mm posterior, ±2.50 mm lateral, 1.85 mm ventral to bregma) and Cre-dependent color-flipping nuclear reporter AAV8-EF1a-Nuc-flox(mCherry)-EGFP^56^ (gift from Brandon Harvey, Addgene, Cat # 112677) was injected in CA1 (1.80 mm posterior, ±1.00 mm lateral, 2.20 mm ventral to bregma). The viral vector carrying a construct coding for the Cre-independent inhibitory DREADD^57^ (AAV8-hSyn-HA-hM4D(Gi)-mCherry, gift from Bryan Roth, Addgene, Cat # 50475) or Cre-dependent inhibitory DREADD^57^ (AAV8-hSyn-DIO-hM4D(Gi)-mCherry, gift from Bryan Roth, Addgene, Cat # 44362) was bilaterally infused into the DH (1.80 mm posterior, ±1.00 mm lateral, 2.25 mm ventral to bregma). For Fiberphotometry experiments viral vector carrying a construct coding for the Cre-dependent GCaMP to DH, CA2 and CA3 area (ssAAV-9/2-hSyn1-chI-dlox-jGCaMP7s(rev)-dlox-WPRE-SV40p(A), VVF Zürich, Cat # v407-9, injected in DH: 1.80 mm posterior, ±1.00 mm lateral, 2.13 mm ventral to bregma and CA3: 1.80 mm posterior, ±1.80 mm lateral, 2.25 mm and ssAAV-5/2-hSyn1-chI-dlox-jGCaMP7s(rev)-dlox-WPRE-SV40p(A), VVF Zürich, Cat # v407-5, injected into CA2: 1.80 mm posterior, ±1.70 mm lateral, 2.05 mm ventral to bregma). For tracing experiment, we injected Cre-dependent viral vector that exobits Synapsin-driven expression of membrane-bound GFP while the mRuby is fused to the Synaptophysin, hence labeling the presynaptic membrane^58^ (gift from Liqun Luo, pAAV1-hSyn-FLEx-mGFP-2A-Synaptophysin-mRuby, Addgene Cat # 71760, CA3: 1.80 mm posterior, ±1.80 mm lateral, 2.25 mm). Using this approach, the cell bodies located at the injection site and their projections express GFP, while mRuby is selectively expressed in regions where the fibers establish synaptic connections.

For behavioral experiments infusions were performed using an automatic microsyringe pump controller (Micro4-WPI) connected to a Hamilton microsyringe (Cat # 88400). The viral vectors were infused in a volume of 0.5 μL per site over 2 min, and syringes were left in place for 5 min prior to removal to allow for virus diffusion. Bilateral 26 gauge guide cannulas (Plastics One) were placed in RSP (1.8 mm posterior, ±0.4 mm lateral, 0.75 mm ventral to bregma). Mice were allowed 6 weeks for virus expression prior to behavioral testing. CNO (Sigma; 0.3 μg/mL; 0.20 μL per side, at a rate of 0.5 μL/min) was infused through the cannulas 30 min prior to either fear conditioning or memory retrieval testing. After the completion of behavioral testing, all brains were collected and cannula placements and virus spread were confirmed by immunohistochemical analysis using anti-mCherry antibodies (1:1000; Abcam, Cat # ab167453). For tracing and fiberphotometry experiments we used Microliter Neuros Syringe (Cat # 65460-02) with an automatic microsyringe pump controller (Micro4-WPI) to deliver unilaterally 0.2 μL of viral vectors over 2 min. Syringes were left in place for 5 min prior to removal to allow for virus diffusion. Optic fibers (MBF Bioscience, Cat # FOC-BF-200-125 with 200 um core diameter and NA=0.37) were implanted to record from SUB→RSP projections (1.80 mm posterior, ±0.30 mm lateral, 1.00 mm ventral to bregma) or 50 μm above injection site for CA2 and CA3 recordings. After the completion of experiments, all brains were collected and fiber placements and virus spread were confirmed by immunohistochemical analysis using anti-GFP antibodies (1:2000; Abcam, Cat # 13970).

### Immunohistochemistry and immunofluorescence

Mice were anesthetized with an i.p. injection of 240 mg/kg Avertin and transcardially perfused with ice-cold 4% paraformaldehyde in phosphate buffer (pH 7.4, 150 mL per mouse). Brains were removed and post-fixed for 48 h in the same fixative and then immersed for 24 h each in 10%, 20% and 30% sucrose solution in phosphate buffer. Brains were frozen and 50 µm sections were cut for use in free-floating immunohistochemistry, as described previously.^59^ Primary antibodies against mCherry (1:1000; Abcam AB167453) and GFP (1:2000, Abcam AB13970) were used and visualized with either diaminobenzidine (Sigma) or secondary antibodies obtained from Jackson ImmunoResearch (1:500 each, AlexaFluor® 594 Cat # 711-585-152, AlexaFluor® 488 Cat # 703-545-155). Sections were mounted using Vectashield (Vector) and observed with a confocal laser-scanning microscope (Olympus Fluoview FV10i) at 40×.

### Quantification of VGluT1^+^ and VGluT2^+^ presynaptic terminal proteomes by *in vivo* proximity biotinylation and TMT-MS

First, we constructed a molecular probe by fusing the promiscuous biotin ligase BirA* to a presynaptic targeting motif (FLEx-preBirA*).^31,32,60^ We also included a T2a motif followed by GFP to visualize neurons expressing our probe and flanking FLEx elements to conditionally express the probe in a cell specific manner based on Cre expression. Once the probe is expressed in the brain, it will localize to presynaptic terminals and biotinylate nearby proteins with a radius of 10 nm. As a negative control, we removed the presynaptic targeting sequence from to generate a construct (cytoFLEx-BirA*) that is not selectively targeted to any subcellular location. Finally, we packaged FLEx-preBirA* and FLEx-cytoBirA* into AAV viruses (packaged by ViroVek). We then sterotactically injected FLEx-preBirA* or FLEx-cytoBirA* AAVs into DH of VGluT1-Cre and VGluT2-Cre mice as described above and incubated the viruses for one month. Next, we administrated biotin (22.5 mg/kg subcutaneously, Sigma B4501) to the mice daily for one week to induce biotinylation of presynaptic proteins. Then we sacrificed the mice and dissected the RSP regions.

Dissected RSPs were homogenized in RIPA lysis buffer (50 mM Tris, 150 mM NaCl, 0.1% SDS, 1mM EDTA, 0.5% sodium deoxycholate, 1% Triton X-100, 1 x protease inhibitor cocktail (Thermo Fisher Scientific, Cat # 78443), 1 x phosphatase inhibitor (Thermo Fisher Scientific, Cat # 78420), pH 7.4) with an electronic homogenizer (Glas-Col, Cat # 099C-K54). Then excess 10% SDS solution was added into each sample to make the final SDS concentration to 1%. After sonication with a probe sonicator (Qsonica) for 3 x 1 min, RSP homogenates were solubilized at 4°C for 1 h with rotation. Insoluble components were removed by centrifuging at 13,000 x g for 30 mins. 400 μL of pre-washed NeutrAvidin beads (Thermo Fisher Scientific, Cat # 2901) were added into each sample and incubated at 4°C overnight with gentle rotation.

We performed on-beads digestion based on previous reported protocol.^61^ After overnight incubation with RSP homogenates, NeutrAvidin beads were rinsed for five times in one mL lysis buffer (6 M Guanidine, 50 mM HEPES, pH 8.5), then added one mL lysis buffer. Dithiothreitol (DTT, DOT Scientific Inc, Cat# DSD11000) was applied to a final concentration of 5 mM. After incubation at RT for 20 min, iodoacetamide (IAA, Sigma-Aldrich, Cat# I1149) was added to a final concentration of 15 mM and incubated for 20 min at room temperature in the dark. Excess IAA was quenched with DTT for 15 min. Samples were diluted with buffer (100 mM HEPES, pH 8.5, 1.5 M Guanidine), and digested for 3 h with Lys-C protease (1:100, ThermoFisher Scientific, Cat# 90307_3668048707) at 37°C. Trypsin (1:100, Promega, Cat# V5280) was then added for overnight incubation at 37°C with intensive agitation (1000 rpm). The next day, reaction was quenched by adding 1% trifluoroacetic acid (TFA, Fisher Scientific, O4902-100). The samples were desalted using HyperSep C18 Cartridges (Thermo Fisher Scientific, Cat# 60108-301) and vacuum centrifuged to dry.

C18 column-desalted peptides were resuspended with 100 mM HEPES pH 8.5 and the concentrations were measured by micro BCA kit (Fisher Scientific, Cat# PI23235). For each sample, 25 μg of peptide labeled with TMT reagent (0.4 mg, dissolved in 40 μL anhydrous acetonitrile, Thermo Fisher Scientific, Cat# 90111) and made at a final concentration of 30% (v/v) acetonitrile (ACN). Following incubation at room temperature for 2 h with agitation, hydroxylamine (to a final concentration of 0.3% (v/v)) was added to quench the reaction for 15 min. Equal amounts of TMT-tagged samples were mixed. Combined sample was vacuum centrifuged to dryness, resuspended, and subjected to HyperSep C18 Cartridges. We used a high pH reverse-phase peptide fractionation kit (Thermo Fisher Scientific, Cat# 84868) to get eight fractions (5.0%, 10.0%, 12.5%, 15.0%, 17.5%, 20.0%, 22.5%, 25.0% and 50% of ACN in 0.1% triethylamine solution). The high pH peptide fractions were directly loaded into the autosampler for MS analysis without further desalting.

3 μg of each fraction or sample were auto-sampler loaded with a Thermo UltiMate 3000 HPLC pump onto a vented Acclaim Pepmap 100, 75 μm x 2 cm, nanoViper trap column coupled to a nanoViper analytical column (Thermo Fisher Scientific, Cat#: 164570, 3 µm, 100 Å, C18, 0.075 mm, 500 mm) with stainless steel emitter tip assembled on the Nanospray Flex Ion Source with a spray voltage of 2000 V. An Orbitrap Fusion (Thermo Fisher Scientific) was used to acquire all the MS spectral data. Buffer A contained 94.785% H_2_O with 5% ACN and 0.125% FA, and buffer B contained 99.875% ACN with 0.125% FA. The chromatographic run was for 4 h in total with the following profile: 0-7% for 7, 10% for 6, 25% for 160, 33% for 40, 50% for 7, 95% for 5 and again 95% for 15 mins receptively.

We used a multiNotch MS3-based TMT method to analyze all the TMT samples ^62–64^. The scan sequence began with an MS1 spectrum (Orbitrap analysis, resolution 120,000, 400-1400 Th, AGC target 2×10^5^, maximum injection time 200 ms). MS2 analysis, ‘Top speed’ (2 s), Collision-induced dissociation (CID, quadrupole ion trap analysis, AGC 4×10^3^, NCE 35, maximum injection time 150 ms). MS3 analysis, top ten precursors, fragmented by HCD prior to Orbitrap analysis (NCE 55, max AGC 5×10^4^, maximum injection time 250 ms, isolation specificity 0.5 Th, resolution 60,000).

Protein identification/quantification and analysis were performed with Integrated Proteomics Pipeline - IP2 (Bruker, Madison, WI. http://www.integratedproteomics.com/) using ProLuCID ^65,66^, DTASelect2 ^67,68^, Census and Quantitative Analysis. Spectrum raw files were extracted into MS1, MS2 and MS3 files using RawConverter (http://fields.scripps.edu/downloads.php). The tandem mass spectra were searched against UniProt mouse protein database (downloaded on 03-25-2014)^69^ and matched to sequences using the ProLuCID/SEQUEST algorithm (ProLuCID version 3.1) with 5 ppm peptide mass tolerance for precursor ions and 600 ppm for fragment ions. The search space included all fully and half-tryptic peptide candidates within the mass tolerance window with no-miscleavage constraint, assembled, and filtered with DTASelect2 through IP2. To estimate peptide probabilities and false-discovery rates (FDR) accurately, we used a target/decoy database containing the reversed sequences of all the proteins appended to the target database.^70^ Each protein identified was required to have a minimum of one peptide of minimal length of six amino acid residues; however, this peptide had to be an excellent match with an FDR < 1% and at least one excellent peptide match. After the peptide/spectrum matches were filtered, we estimated that the peptide FDRs were ≤ 1% for each sample analysis. Resulting protein lists include subset proteins to allow for consideration of all possible protein forms implicated by at least two given peptides identified from the complex protein mixtures. Then, we used Census and Quantitative Analysis in IP2 for protein quantification of TMT MS. experiments and protein quantification was determined by summing all TMT report ion counts. TMT MS data were normalized using with a build-in method in IP2.

Spyder (MIT, Python 3.7, libraries, ‘pandas’, ‘numpy’, ‘scipy’, ‘statsmodels’ and ‘bioinfokit’) was used for data analyses. RStudio (version, 1.2.1335, packages, ‘tidyverse’, ‘pheatmap’) was used for data virtualization. The Database for Annotation, Visualization and Integrated Discovery (DAVID) (https://david.ncifcrf.gov/) was used for protein functional annotation analysis.

### Single cell RNA sequencing

The snRNA seq data shown in this publication was obtained by analyzing dataset previously deposited to NCBI’s Gene Expression Omnibus by our group^33^ and can be accessed using GEO Series accession number GSE254780 (https://www.ncbi.nlm.nih.gov/geo/query/acc.cgi?acc=GSE254780). Briefly, single-cell RNA sequencing (scRNA-seq) data were initially analyzed using scTE with default settings to quantify transposable element (TE) gene expression.^71^ The resulting .h5ad data objects were loaded into R (v4.3) and analyzed using the Seurat package (v5.0.3).^72^ Within Seurat, cells were filtered to retain those with >500 UMI counts, >200 detected features, and <15% mitochondrial content.

Genes expressed in fewer than 5 cells were also removed. Doublets were subsequently identified and removed using DoubletFinder.^73^ The filtered datasets were then merged. Principal Component Analysis (PCA) was performed, and an optimal number of PCs was selected based on cumulative variance and elbow point heuristics. Batch effects between samples were corrected using Harmony integration on the PCA embeddings.^74^ UMAP visualization and Leiden clustering (resolution 0.4) were performed on the Harmony-corrected dimensions. Cell type annotation was performed using scType with a custom brain-specific marker gene database (Supplementary table 2).^75^ Finally, differentially expressed genes (DEGs) between annotated cell types were identified using FindAllMarkers function from Seurat package, and these DEGs were further filtered to identify transposable element (TE)-derived transcripts based on a provided TE annotation file provided by the scTE package. The expression of VGluT2 (*Slc17a6*) and VGlut1 (*Slc17a7*) genes, was examined using FeaturePlot function from the Seurat package with the blend option set to TRUE, enabling the visualization of co-expression patterns between these genes. Data showing expression of VGluT2 (*Slc17a6*) and VGlut1 (*Slc17a7*) genes in the subicular clusters was generated using database previously published^28^ and available for online analysis on Brain RNA-seq atlas (https://scrnaseq.janelia.org/). Following online analysis with the available interactive tool, gene expression values were downloaded and plotted offline using GraphPad Prizm software.

### Trace fear conditioning

Trace fear conditioning was performed in an automated system (TSE Systems) as described previously. ^76^ Briefly, mice were exposed for 120 s to a unfamiliar context (Context 1), followed by a 30 s of 10 kHz tone, 15 s temporal trace, and foot shock (2 s, 0.7 mA, constant current). To prevent scent-based cues from influencing behavior, chambers were cleaned thoroughly with 70% ethanol after every session. Mice were tested for memory retrieval 24 h later in a contextually distinct novel context (Context 2, context exposure duration 60 s, Tone duration 30 s and Trace duration 15 s). Chambers were cleaned thoroughly with 1% acetic acid after every session. Freezing was scored every 5 s during Context 2 and Tone exposure and every 3 s during Trace duration, and expressed as a percentage of the total number of freezing observations during which the mice were motionless. To control for the behavioral specificity of CNO effects, we also used delay fear conditioning (DFC) and tone-light-shock fear conditioning (TLC). DFC was performed as TFC except that shock was delivered immediately after termination of tone, thus there was no trace separating tone and shock. TLC was performed as TFC except that 500 ms light pulses were delivered during the 15 s trace. For fiberphotometry experiments involving trace fear conditioning, mice were placed in soundproof chambers (Med Associates Inc., St. Albans, VT) equipped with metal-grid flooring used to administer foot shocks. Each session began with a 120 s of context exposure (Context 1), after which the animals were presented with three auditory tones (30 s duration, 80 dB, 50 ms rise time). A brief foot shock (0.5 mA, 2 s) was delivered 15 s after each tone ended. Each Tone-Trace presentation was separated by a 60 s interval. To prevent scent-based cues from influencing behavior, chambers were cleaned thoroughly with 70% ethanol after every session. Memory recall was evaluated 24 hours later in a contextually distinct environment (Context 2) featuring a flat white floor and a novel odor (1% acetic acid). Mice were placed in this altered chamber and, following a 120 s baseline period, the tone was played again for 30 s. Freezing responses were measured during the Baseline, Tone, and Trace period. Behavior was recorded and quantified using Video Freeze® software (Med Associates Inc., Fairfax, VT), and independently validated by manual scoring (Context 2 and Tone exposure every 5 s and Trace every 3 s) by experimenters blinded to both treatment groups and experimental design through the use of coded identifiers. All behavioral experiments were performed between 10 am and 5 pm. Littermates were randomly assigned to the different treatment conditions. All behavioral tests and immunohistochemical analyses were performed by experimenters who were blind to genotypes and drug treatments.

### Fiberphotometry

We used fiberphotomerty technique to measure real time neuronal calcium transients in freely moving animals.^77,78^ Data acquisition was performed using Neurophotometrics FP3002 system using 2 LEDs with emission of 470 nm (GCaMP wavelength) and 415 nm (isosbestic wavelength) coupled to a 200 μm 0.37 N.A. optical fiber (FOC-BF-200-125, Neurophotmetrics) with the light power at the tip constant across trials and testing days between 10-30 μW. The fluorescence signal was collected by the same optical fiber, filtered, and focused on a BlackFly CMOS camera. Intercalated samples were acquired at 60 FPS. Raw data were used as an input and visualized by Bonsai^79^ and CropPolygon function was used to define ROIs. Synchronization with movement was provided by simultaneously triggering a TTL input from the fear conditioning system (MedAssociates Inc., St. Albans, VT). To minimize the patch cord autofluorescence, prior to the recording light at 470 nm was delivered for at least 15 h. Animals were handled and habituated in their home cage to optical fiber tethering for 3 days prior to behavioral tests.

For data analysis we used GuPPy, aPython toolbox for fiberphotometry analysis.^80^ The isosbestic wavelength records calcium-independent events such as motion artifacts and autofluorescence, and photobleaches at the similar rate as the calcium signal and was used as a control signal when calculating fluorescence signal changes from the baseline, using following equation ΔF/F =(F_observed_-F_fitted_)/F_fitted_.^81^ High-pass filtered and transformed to z scores data (z score = (ΔF/F-μ_ΔF/F_))/σ_ΔF/F_, where μ is mean and σ is standard deviation) was used to combine data across multiple animals and testing days. Ca^2+^ activity associated with different test phases was assessed by aligning the ΔF/F signal to time 0 at each TTL timestamp and extracting a 90 s window, spanning 30 s before to 60 s after Tone onset. Data was binned to 15 s intervals and positive and negative areas under the curve (AUC)^82^ were calculated for each detected peaks and average z score for each training and testing phase.

### Quantification and statistical analyses

Statistical analyses were performed using Graphpad software and Matlab functions. For the behavioral studies, freezing data were analyzed for Treatment (CNO or Vehicle) and Test (repeated measure) as factors using two-way repeated measures ANOVA. For fiber photometry studies two-way repeated measures ANOVA was used to compare between phases during training and testing days (Baseline, Tone, Trace, Inter-Trial-Interval). Significant *F* values were followed by *post hoc* comparisons using Tukey test. For analyses of proteomic data, we used regression analyses and unpaired two-tailed Student’s *t* test. Homogeneity of variance was confirmed with Levene’s test for equality of variances. Statistical differences were considered significant for all *P* values < 0.05. Group sizes were determined using power analyses assuming a moderate effect size of 0.5. All key findings were replicated at least twice, and mostly three times, in different sets of mice (biological replicates). Only mice with correctly placed cannulas and fibers and robust virus expression in RSP terminals or injection site (> 70% of maximal expression determined by densitometry) were included in the analyses. Details of statistical analyses are found in figure legends. All data for the preparation of graphs and statistical analysis relevant data that support the conclusions are uploaded as source data.

## References

1. Tulving, E. (1972). Episodic and semantic memory. Organization of memory 1, 1.

2. Tulving, E. (1985). Memory and consciousness. Canadian Psychology/Psychologie canadienne 26, 1.

3. Eichenbaum, H. (2014). Time cells in the hippocampus: a new dimension for mapping memories. Nat Rev Neurosci 15, 732–744. 10.1038/nrn3827.

4. Kitamura, T. (2017). Driving and regulating temporal association learning coordinated by entorhinal-hippocampal network. Neurosci Res 121, 1–6. 10.1016/j.neures.2017.04.005.

5. Kim, J.J., DeCola, J.P., Landeira-Fernandez, J., and Fanselow, M.S. (1991). N-methyl-D-aspartate receptor antagonist APV blocks acquisition but not expression of fear conditioning. Behav Neurosci 105, 126–133. 10.1037//0735-7044.105.1.126.

6. Misane, I., Tovote, P., Meyer, M., Spiess, J., Ogren, S.O., and Stiedl, O. (2005). Time-dependent involvement of the dorsal hippocampus in trace fear conditioning in mice. Hippocampus 15, 418–426. 10.1002/hipo.20067.

7. Gilmartin, M.R., Miyawaki, H., Helmstetter, F.J., and Diba, K. (2013). Prefrontal activity links nonoverlapping events in memory. J Neurosci 33, 10910–10914. 10.1523/JNEUROSCI.0144-13.2013.

8. Huerta, P.T., Sun, L.D., Wilson, M.A., and Tonegawa, S. (2000). Formation of temporal memory requires NMDA receptors within CA1 pyramidal neurons. Neuron 25, 473–480.

9. Puhger, K., Crestani, A.P., Diniz, C., and Wiltgen, B.J. (2024). The hippocampus contributes to retroactive stimulus associations during trace fear conditioning. iScience 27, 109035. 10.1016/j.isci.2024.109035.

10. Roy, D.S., Kitamura, T., Okuyama, T., Ogawa, S.K., Sun, C., Obata, Y., Yoshiki, A., and Tonegawa, S. (2017). Distinct Neural Circuits for the Formation and Retrieval of Episodic Memories. Cell 170, 1000–1012 e1019. 10.1016/j.cell.2017.07.013.

11. Suh, J., Rivest, A.J., Nakashiba, T., Tominaga, T., and Tonegawa, S. (2011). Entorhinal cortex layer III input to the hippocampus is crucial for temporal association memory. Science 334, 1415–1420.

12. Kitamura, T., Pignatelli, M., Suh, J., Kohara, K., Yoshiki, A., Abe, K., and Tonegawa, S. (2014). Island cells control temporal association memory. Science 343, 896–901.

13. Bai, T., Zhan, L., Zhang, N., Lin, F., Saur, D., and Xu, C. (2023). Learning-prolonged maintenance of stimulus information in CA1 and subiculum during trace fear conditioning. Cell Rep 42, 112853. 10.1016/j.celrep.2023.112853.

14. Kwapis, J.L., Jarome, T.J., Lee, J.L., Gilmartin, M.R., and Helmstetter, F.J. (2014). Extinguishing trace fear engages the retrosplenial cortex rather than the amygdala. Neurobiology of learning and memory 113, 41–54.

15. Kwapis, J.L., Jarome, T.J., Lee, J.L., and Helmstetter, F.J. (2015). The retrosplenial cortex is involved in the formation of memory for context and trace fear conditioning. Neurobiology of Learning and Memory 123, 110–116.

16. Opalka, A.N., and Wang, D.V. (2020). Hippocampal efferents to retrosplenial cortex and lateral septum are required for memory acquisition. Learn Mem 27, 310–318. 10.1101/lm.051797.120.

17. Garvert, A.C., Bieler, M., Witoelar, A., and Vervaeke, K. (2025). Area-specific encoding of temporal information in the neocortex. Cell Rep 44, 115363. 10.1016/j.celrep.2025.115363.

18. Yamawaki, N., Li, X., Lambot, L., Ren, L.Y., Radulovic, J., and Shepherd, G.M.G. (2019). Long-range inhibitory intersection of a retrosplenial thalamocortical circuit by apical tuft-targeting CA1 neurons. Nat Neurosci 22, 618–626. 10.1038/s41593-019-0355-x.

19. Yamawaki, N., Corcoran, K.A., Guedea, A.L., Shepherd, G.M.G., and Radulovic, J. (2019). Differential Contributions of Glutamatergic Hippocampal-->Retrosplenial Cortical Projections to the Formation and Persistence of Context Memories. Cereb Cortex 29, 2728–2736. 10.1093/cercor/bhy142.

20. Corcoran, K.A., Yamawaki, N., Leaderbrand, K., and Radulovic, J. (2018). Role of retrosplenial cortex in processing stress-related context memories. Behav Neurosci 132, 388–395. 10.1037/bne0000223.

21. McEchron, M.D., Bouwmeester, H., Tseng, W., Weiss, C., and Disterhoft, J.F. (1998). Hippocampectomy disrupts auditory trace fear conditioning and contextual fear conditioning in the rat. Hippocampus 8, 638–646. 10.1002/(SICI)1098-1063(1998)8:6<638::AID-HIPO6>3.0.CO;2-Q.

22. Kwapis, J.L., Jarome, T.J., Lee, J.L., and Helmstetter, F.J. (2015). The retrosplenial cortex is involved in the formation of memory for context and trace fear conditioning. Neurobiol Learn Mem 123, 110–116. 10.1016/j.nlm.2015.06.007.

23. Stachniak, T.J., Ghosh, A., and Sternson, S.M. (2014). Chemogenetic synaptic silencing of neural circuits localizes a hypothalamus-->midbrain pathway for feeding behavior. Neuron 82, 797–808. 10.1016/j.neuron.2014.04.008.

24. Harris, J.A., Hirokawa, K.E., Sorensen, S.A., Gu, H., Mills, M., Ng, L.L., Bohn, P., Mortrud, M., Ouellette, B., Kidney, J., et al. (2014). Anatomical characterization of Cre driver mice for neural circuit mapping and manipulation. Front Neural Circuits 8, 76. 10.3389/fncir.2014.00076.

25. Vong, L., Ye, C., Yang, Z., Choi, B., Chua, S., Jr., and Lowell, B.B. (2011). Leptin action on GABAergic neurons prevents obesity and reduces inhibitory tone to POMC neurons. Neuron 71, 142–154. 10.1016/j.neuron.2011.05.028.

26. El Mestikawy, S., Wallen-Mackenzie, A., Fortin, G.M., Descarries, L., and Trudeau, L.E. (2011). From glutamate co-release to vesicular synergy: vesicular glutamate transporters. Nat Rev Neurosci 12, 204–216. 10.1038/nrn2969.

27. Fremeau, R.T., Jr., Troyer, M.D., Pahner, I., Nygaard, G.O., Tran, C.H., Reimer, R.J., Bellocchio, E.E., Fortin, D., Storm-Mathisen, J., and Edwards, R.H. (2001). The expression of vesicular glutamate transporters defines two classes of excitatory synapse. Neuron 31, 247–260. 10.1016/s0896-6273(01)00344-0.

28. Cembrowski, M.S., Wang, L., Lemire, A.L., Copeland, M., DiLisio, S.F., Clements, J., and Spruston, N. (2018). The subiculum is a patchwork of discrete subregions. Elife 7. 10.7554/eLife.37701.

29. Cembrowski, M.S., Wang, L., Sugino, K., Shields, B.C., and Spruston, N. (2016). Hipposeq: a comprehensive RNA-seq database of gene expression in hippocampal principal neurons. Elife 5, e14997. 10.7554/eLife.14997.

30. Cembrowski, M.S., Phillips, M.G., DiLisio, S.F., Shields, B.C., Winnubst, J., Chandrashekar, J., Bas, E., and Spruston, N. (2018). Dissociable Structural and Functional Hippocampal Outputs via Distinct Subiculum Cell Classes. Cell 174, 1036. 10.1016/j.cell.2018.07.039.

31. Wang, Y.Z., Perez-Rosello, T., Smukowski, S.N., Surmeier, D.J., and Savas, J.N. (2024). Neuron type-specific proteomics reveals distinct Shank3 proteoforms in iSPNs and dSPNs lead to striatal synaptopathy in Shank3B(-/-) mice. Mol Psychiatry 29, 2372–2388. 10.1038/s41380-024-02493-w.

32. Kim, J., Zhao, T., Petralia, R.S., Yu, Y., Peng, H., Myers, E., and Magee, J.C. (2011). mGRASP enables mapping mammalian synaptic connectivity with light microscopy. Nat Methods 9, 96–102. 10.1038/nmeth.1784.

33. Jovasevic, V., Wood, E.M., Cicvaric, A., Zhang, H., Petrovic, Z., Carboncino, A., Parker, K.K., Bassett, T.E., Moltesen, M., Yamawaki, N., et al. (2024). Formation of memory assemblies through the DNA-sensing TLR9 pathway. Nature 628, 145–153. 10.1038/s41586-024-07220-7.

34. Curzon, P., Rustay, N.R., and Browman, K.E. (2009). Cued and Contextual Fear Conditioning for Rodents. In Methods of Behavior Analysis in Neuroscience, J.J. Buccafusco, ed.

35. Berry, C.T., Sceniak, M.P., Zhou, L., and Sabo, S.L. (2012). Developmental up-regulation of vesicular glutamate transporter-1 promotes neocortical presynaptic terminal development. PLoS One 7, e50911. 10.1371/journal.pone.0050911.

36. Guillaud, L., Dimitrov, D., and Takahashi, T. (2017). Presynaptic morphology and vesicular composition determine vesicle dynamics in mouse central synapses. Elife 6. 10.7554/eLife.24845.

37. Weston, M.C., Nehring, R.B., Wojcik, S.M., and Rosenmund, C. (2011). Interplay between VGLUT isoforms and endophilin A1 regulates neurotransmitter release and short-term plasticity. Neuron 69, 1147–1159. 10.1016/j.neuron.2011.02.002.

38. Cavalieri, D., Angelova, A., Islah, A., Lopez, C., Bocchio, M., Bollmann, Y., Baude, A., and Cossart, R. (2021). CA1 pyramidal cell diversity is rooted in the time of neurogenesis. Elife 10. 10.7554/eLife.69270.

39. Mizuseki, K., Diba, K., Pastalkova, E., and Buzsaki, G. (2011). Hippocampal CA1 pyramidal cells form functionally distinct sublayers. Nat Neurosci 14, 1174–1181. 10.1038/nn.2894.

40. Pilkiw, M., and Takehara-Nishiuchi, K. (2018). Neural representations of time-linked memory. Neurobiol Learn Mem 153, 57–70. 10.1016/j.nlm.2018.03.024.

41. Raybuck, J.D., and Lattal, K.M. (2014). Bridging the interval: theory and neurobiology of trace conditioning. Behav Processes 101, 103–111. 10.1016/j.beproc.2013.08.016.

42. Veres, J.M., Andrasi, T., Nagy-Pal, P., and Hajos, N. (2023). CaMKIIalpha Promoter-Controlled Circuit Manipulations Target Both Pyramidal Cells and Inhibitory Interneurons in Cortical Networks. eNeuro 10. 10.1523/ENEURO.0070-23.2023.

43. Ahmed, M.S., Priestley, J.B., Castro, A., Stefanini, F., Solis Canales, A.S., Balough, E.M., Lavoie, E., Mazzucato, L., Fusi, S., and Losonczy, A. (2020). Hippocampal Network Reorganization Underlies the Formation of a Temporal Association Memory. Neuron 107, 283–291 e286. 10.1016/j.neuron.2020.04.013.

44. Gilmartin, M.R., and McEchron, M.D. (2005). Single neurons in the dentate gyrus and CA1 of the hippocampus exhibit inverse patterns of encoding during trace fear conditioning. Behav Neurosci 119, 164–179. 10.1037/0735-7044.119.1.164.

45. Jinno, S., Klausberger, T., Marton, L.F., Dalezios, Y., Roberts, J.D., Fuentealba, P., Bushong, E.A., Henze, D., Buzsaki, G., and Somogyi, P. (2007). Neuronal diversity in GABAergic long-range projections from the hippocampus. J Neurosci 27, 8790–8804. 10.1523/JNEUROSCI.1847-07.2007.

46. Csicsvari, J., Hirase, H., Mamiya, A., and Buzsaki, G. (2000). Ensemble patterns of hippocampal CA3-CA1 neurons during sharp wave-associated population events. Neuron 28, 585–594. 10.1016/s0896-6273(00)00135-5.

47. Imbrosci, B., Nitzan, N., McKenzie, S., Donoso, J.R., Swaminathan, A., Bohm, C., Maier, N., and Schmitz, D. (2021). Subiculum as a generator of sharp wave-ripples in the rodent hippocampus. Cell Rep 35, 109021. 10.1016/j.celrep.2021.109021.

48. Oliva, A., Fernandez-Ruiz, A., Buzsaki, G., and Berenyi, A. (2016). Role of Hippocampal CA2 Region in Triggering Sharp-Wave Ripples. Neuron 91, 1342–1355. 10.1016/j.neuron.2016.08.008.

49. Kitanishi, T., Umaba, R., and Mizuseki, K. (2021). Robust information routing by dorsal subiculum neurons. Sci Adv 7. 10.1126/sciadv.abf1913.

50. Picard, L., Cousin, S., Guillery-Girard, B., Eustache, F., and Piolino, P. (2012). How do the different components of episodic memory develop? Role of executive functions and short-term feature-binding abilities. Child Dev 83, 1037–1050. 10.1111/j.1467-8624.2012.01736.x.

51. Lee, J.K., Wendelken, C., Bunge, S.A., and Ghetti, S. (2016). A Time and Place for Everything: Developmental Differences in the Building Blocks of Episodic Memory. Child Dev 87, 194–210. 10.1111/cdev.12447.

52. Hitti, F.L., and Siegelbaum, S.A. (2014). The hippocampal CA2 region is essential for social memory. Nature 508, 88–92. 10.1038/nature13028.

53. Nakazawa, K., Quirk, M.C., Chitwood, R.A., Watanabe, M., Yeckel, M.F., Sun, L.D., Kato, A., Carr, C.A., Johnston, D., Wilson, M.A., and Tonegawa, S. (2002). Requirement for hippocampal CA3 NMDA receptors in associative memory recall. Science 297, 211–218. 10.1126/science.1071795.

54. Yamawaki, N., Corcoran, K.A., Guedea, A.L., Shepherd, G.M.G., and Radulovic, J. (2018). Differential Contributions of Glutamatergic Hippocampal-->Retrosplenial Cortical Projections to the Formation and Persistence of Context Memories. Cereb Cortex. 10.1093/cercor/bhy142.

55. Saunders, A., Johnson, C.A., and Sabatini, B.L. (2012). Novel recombinant adeno-associated viruses for Cre activated and inactivated transgene expression in neurons. Front Neural Circuits 6, 47. 10.3389/fncir.2012.00047.

56. Back, S., Necarsulmer, J., Whitaker, L.R., Coke, L.M., Koivula, P., Heathward, E.J., Fortuno, L.V., Zhang, Y., Yeh, C.G., Baldwin, H.A., et al. (2019). Neuron-Specific Genome Modification in the Adult Rat Brain Using CRISPR-Cas9 Transgenic Rats. Neuron 102, 105–119 e108. 10.1016/j.neuron.2019.01.035.

57. Krashes, M.J., Koda, S., Ye, C., Rogan, S.C., Adams, A.C., Cusher, D.S., Maratos-Flier, E., Roth, B.L., and Lowell, B.B. (2011). Rapid, reversible activation of AgRP neurons drives feeding behavior in mice. J Clin Invest 121, 1424–1428. 10.1172/JCI46229.

58. Beier, K.T., Steinberg, E.E., DeLoach, K.E., Xie, S., Miyamichi, K., Schwarz, L., Gao, X.J., Kremer, E.J., Malenka, R.C., and Luo, L. (2015). Circuit Architecture of VTA Dopamine Neurons Revealed by Systematic Input-Output Mapping. Cell 162, 622–634. 10.1016/j.cell.2015.07.015.

59. Jovasevic, V., Corcoran, K.A., Leaderbrand, K., Yamawaki, N., Guedea, A.L., Chen, H.J., Shepherd, G.M., and Radulovic, J. (2015). GABAergic mechanisms regulated by miR-33 encode state-dependent fear. Nat Neurosci 18, 1265–1271. 10.1038/nn.4084.

60. Wang, Y.Z., and Savas, J.N. (2018). Uncovering Discrete Synaptic Proteomes to Understand Neurological Disorders. Proteomes 6. 10.3390/proteomes6030030.

61. Loh, K.H., Stawski, P.S., Draycott, A.S., Udeshi, N.D., Lehrman, E.K., Wilton, D.K., Svinkina, T., Deerinck, T.J., Ellisman, M.H., Stevens, B., et al. (2016). Proteomic Analysis of Unbounded Cellular Compartments: Synaptic Clefts. Cell 166, 1295–1307 e1221. 10.1016/j.cell.2016.07.041.

62. McAlister, G.C., Nusinow, D.P., Jedrychowski, M.P., Wuhr, M., Huttlin, E.L., Erickson, B.K., Rad, R., Haas, W., and Gygi, S.P. (2014). MultiNotch MS3 enables accurate, sensitive, and multiplexed detection of differential expression across cancer cell line proteomes. Anal Chem 86, 7150–7158. 10.1021/ac502040v.

63. Ting, L., Rad, R., Gygi, S.P., and Haas, W. (2011). MS3 eliminates ratio distortion in isobaric multiplexed quantitative proteomics. Nat Methods 8, 937–940. 10.1038/nmeth.1714.

64. Weekes, M.P., Tomasec, P., Huttlin, E.L., Fielding, C.A., Nusinow, D., Stanton, R.J., Wang, E.C., Aicheler, R., Murrell, I., Wilkinson, G.W., et al. (2014). Quantitative temporal viromics: an approach to investigate host-pathogen interaction. Cell 157, 1460–1472. 10.1016/j.cell.2014.04.028.

65. Eng, J.K., McCormack, A.L., and Yates, J.R. (1994). An approach to correlate tandem mass spectral data of peptides with amino acid sequences in a protein database. J Am Soc Mass Spectrom 5, 976–989. 10.1016/1044-0305(94)80016-2.

66. Xu, T., Park, S.K., Venable, J.D., Wohlschlegel, J.A., Diedrich, J.K., Cociorva, D., Lu, B., Liao, L., Hewel, J., Han, X., et al. (2015). ProLuCID: An improved SEQUEST-like algorithm with enhanced sensitivity and specificity. J Proteomics 129, 16–24. 10.1016/j.jprot.2015.07.001.

67. Cociorva, D., D, L.T., and Yates, J.R. (2007). Validation of tandem mass spectrometry database search results using DTASelect. Curr Protoc Bioinformatics Chapter 13, Unit 13 14. 10.1002/0471250953.bi1304s16.

68. Tabb, D.L., McDonald, W.H., and Yates, J.R., 3rd (2002). DTASelect and Contrast: tools for assembling and comparing protein identifications from shotgun proteomics. J Proteome Res 1, 21–26.

69. UniProt, C. (2015). UniProt: a hub for protein information. Nucleic Acids Res 43, D204–212. 10.1093/nar/gku989.

70. Peng, J., Elias, J.E., Thoreen, C.C., Licklider, L.J., and Gygi, S.P. (2003). Evaluation of multidimensional chromatography coupled with tandem mass spectrometry (LC/LC-MS/MS) for large-scale protein analysis: the yeast proteome. J Proteome Res 2, 43–50.

71. He, J., Babarinde, I.A., Sun, L., Xu, S., Chen, R., Shi, J., Wei, Y., Li, Y., Ma, G., Zhuang, Q., et al. (2021). Identifying transposable element expression dynamics and heterogeneity during development at the single-cell level with a processing pipeline scTE. Nature Communications 12, 1456. 10.1038/s41467-021-21808-x.

72. Hao, Y., Stuart, T., Kowalski, M.H., Choudhary, S., Hoffman, P., Hartman, A., Srivastava, A., Molla, G., Madad, S., and Fernandez-Granda, C. (2024). Dictionary learning for integrative, multimodal and scalable single-cell analysis. Nature biotechnology 42, 293–304.

73. McGinnis, C.S., Murrow, L.M., and Gartner, Z.J. (2019). DoubletFinder: Doublet Detection in Single-Cell RNA Sequencing Data Using Artificial Nearest Neighbors. Cell Systems 8, 329–337.e324. 10.1016/j.cels.2019.03.003.

74. Korsunsky, I., Millard, N., Fan, J., Slowikowski, K., Zhang, F., Wei, K., Baglaenko, Y., Brenner, M., Loh, P.-r., and Raychaudhuri, S. (2019). Fast, sensitive and accurate integration of single-cell data with Harmony. Nature methods 16, 1289–1296.

75. Ianevski, A., Giri, A.K., and Aittokallio, T. (2022). Fully-automated and ultra-fast cell-type identification using specific marker combinations from single-cell transcriptomic data. Nature Communications 13, 1246. 10.1038/s41467-022-28803-w.

76. Corcoran, K.A., Donnan, M.D., Tronson, N.C., Guzman, Y.F., Gao, C., Jovasevic, V., Guedea, A.L., and Radulovic, J. (2011). NMDA receptors in retrosplenial cortex are necessary for retrieval of recent and remote context fear memory. J Neurosci 31, 11655–11659. 10.1523/JNEUROSCI.2107-11.201131/32/11655 [pii].

77. Gunaydin, L.A., Grosenick, L., Finkelstein, J.C., Kauvar, I.V., Fenno, L.E., Adhikari, A., Lammel, S., Mirzabekov, J.J., Airan, R.D., Zalocusky, K.A., et al. (2014). Natural neural projection dynamics underlying social behavior. Cell 157, 1535–1551. 10.1016/j.cell.2014.05.017.

78. Martianova, E., Aronson, S., and Proulx, C.D. (2019). Multi-Fiber Photometry to Record Neural Activity in Freely-Moving Animals. J Vis Exp. 10.3791/60278.

79. Lopes, G., Bonacchi, N., Frazao, J., Neto, J.P., Atallah, B.V., Soares, S., Moreira, L., Matias, S., Itskov, P.M., Correia, P.A., et al. (2015). Bonsai: an event-based framework for processing and controlling data streams. Front Neuroinform 9, 7. 10.3389/fninf.2015.00007.

80. Sherathiya, V.N., Schaid, M.D., Seiler, J.L., Lopez, G.C., and Lerner, T.N. (2021). GuPPy, a Python toolbox for the analysis of fiber photometry data. Sci Rep 11, 24212. 10.1038/s41598-021-03626-9.

81. Lerner, T.N., Shilyansky, C., Davidson, T.J., Evans, K.E., Beier, K.T., Zalocusky, K.A., Crow, A.K., Malenka, R.C., Luo, L., Tomer, R., and Deisseroth, K. (2015). Intact-Brain Analyses Reveal Distinct Information Carried by SNc Dopamine Subcircuits. Cell 162, 635–647. 10.1016/j.cell.2015.07.014.

82. Holly, E.N., Davatolhagh, M.F., Choi, K., Alabi, O.O., Vargas Cifuentes, L., and Fuccillo, M.V. (2019). Striatal Low-Threshold Spiking Interneurons Regulate Goal-Directed Learning. Neuron 103, 92–101 e106. 10.1016/j.neuron.2019.04.016.

