## Supplementary figures and images for "Response dynamics of discrete subiculum→retrosplenial cortex projections underlying trace fear conditioning"

### Figure S1

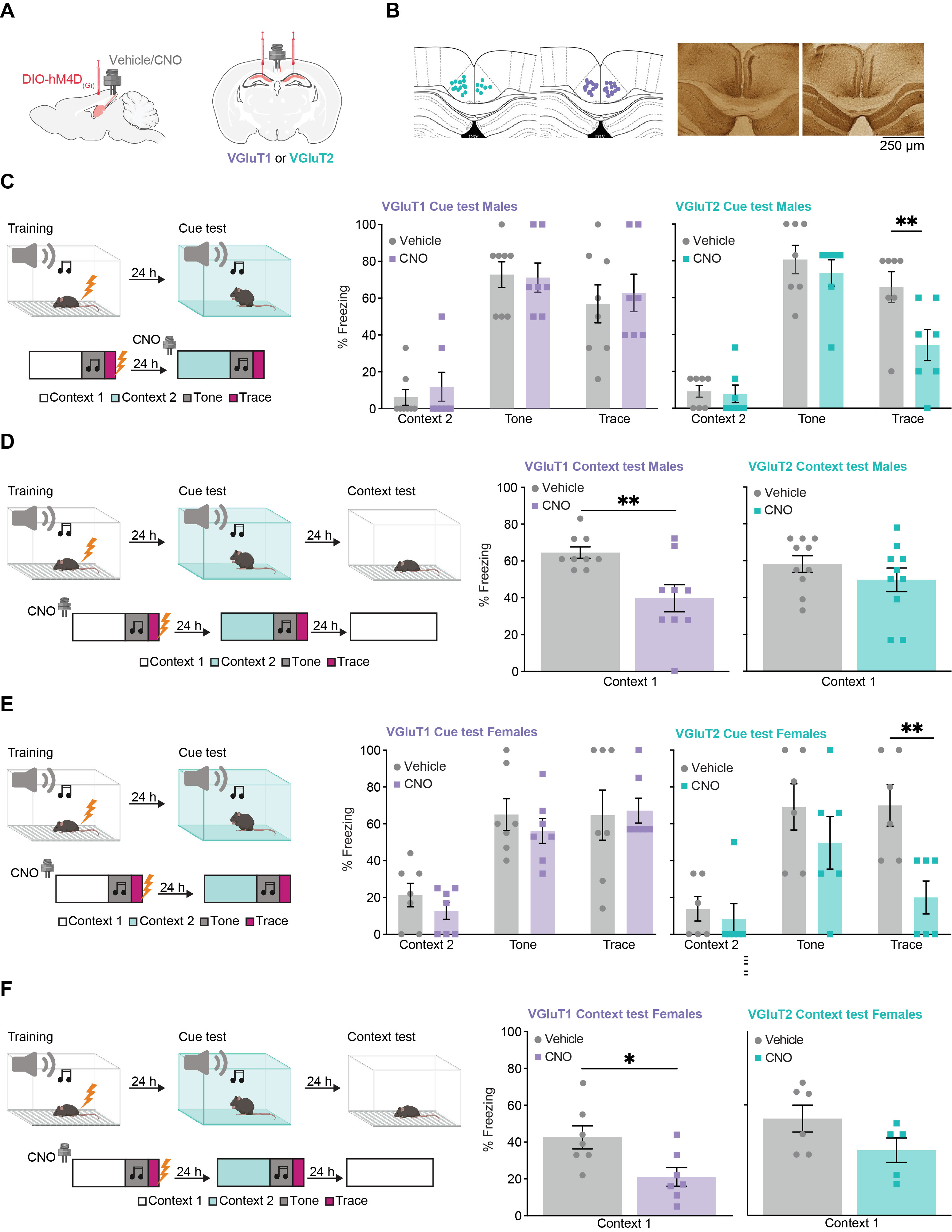

### Figure S2

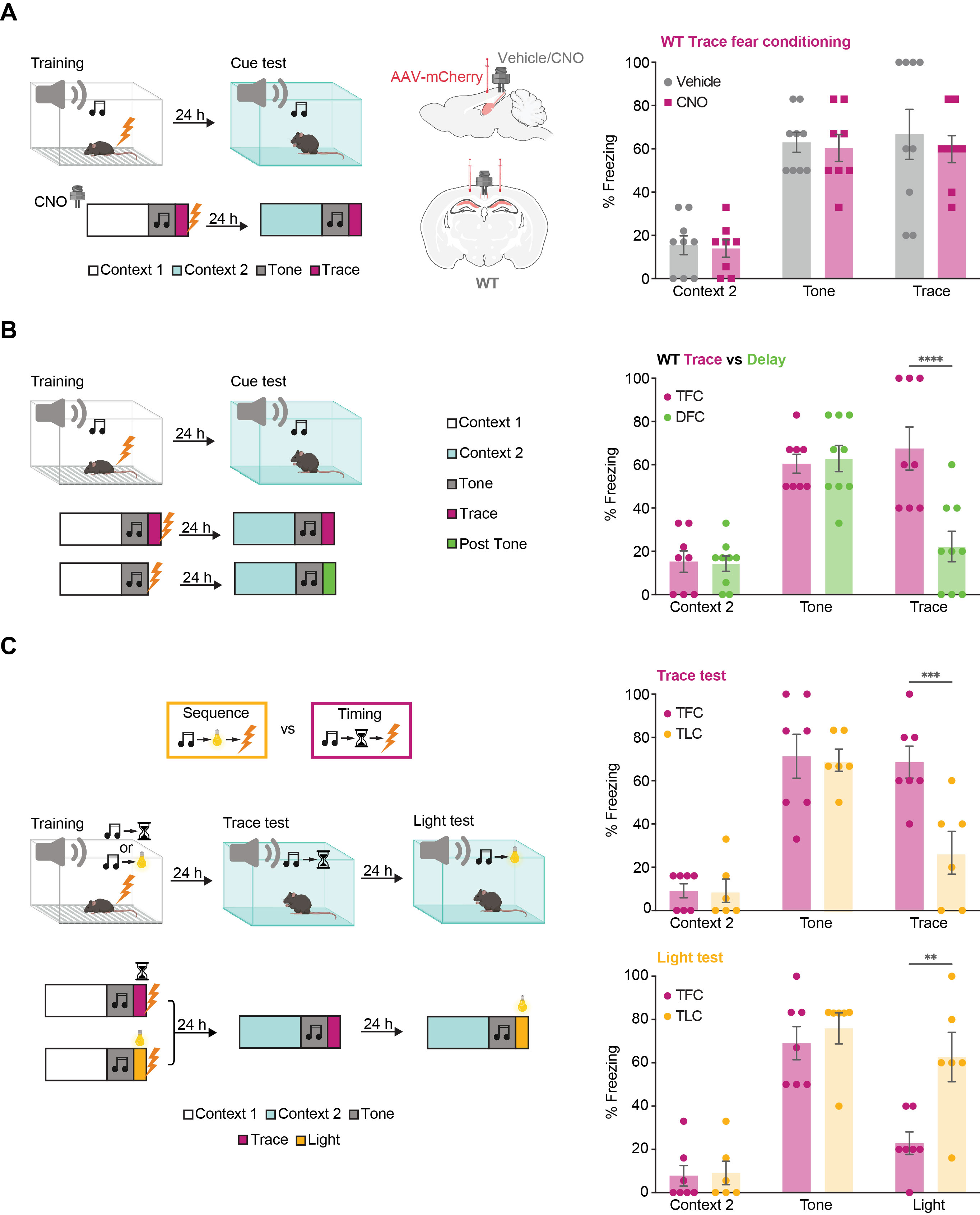

### Figure S3

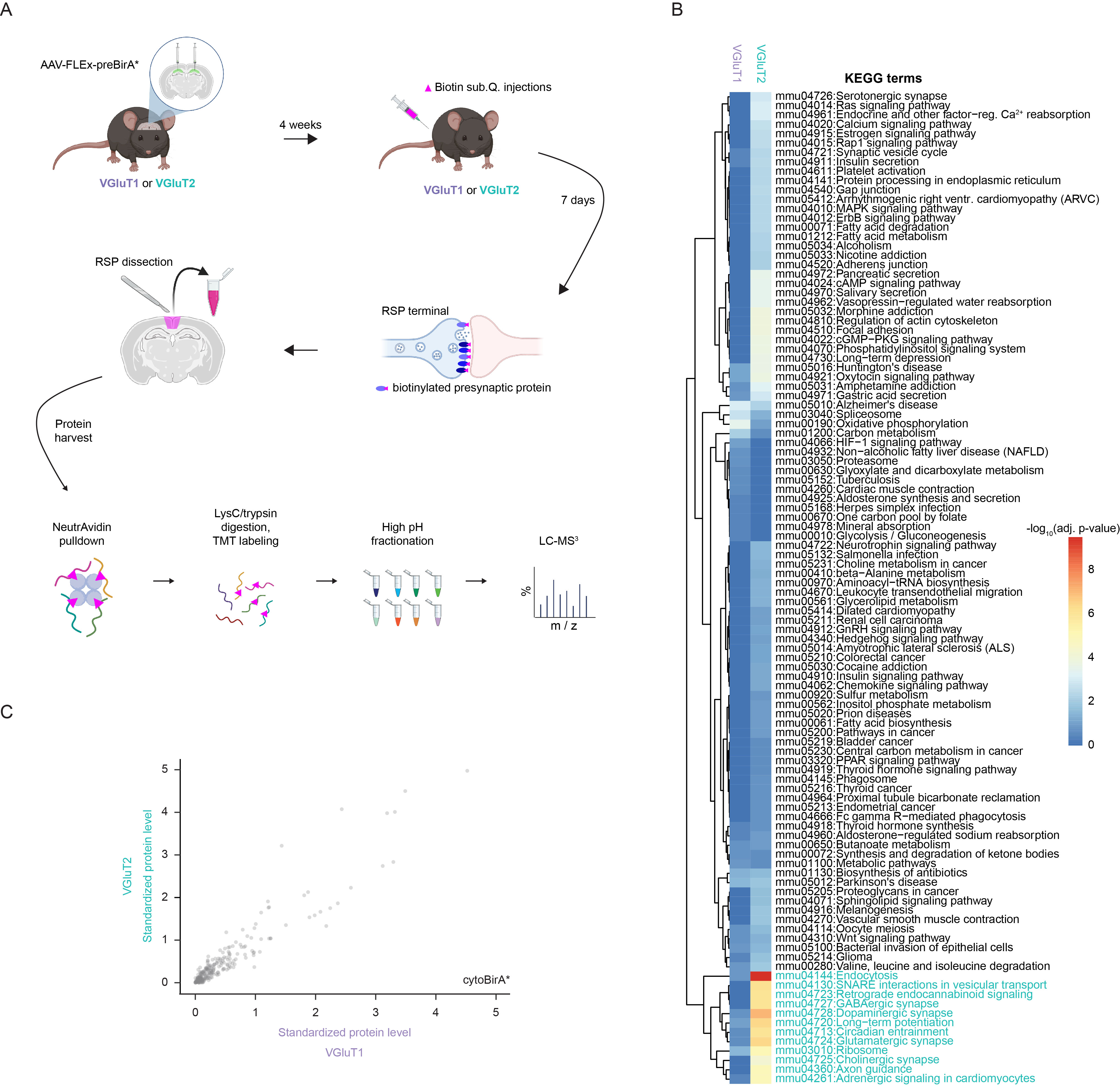

### Figure S4

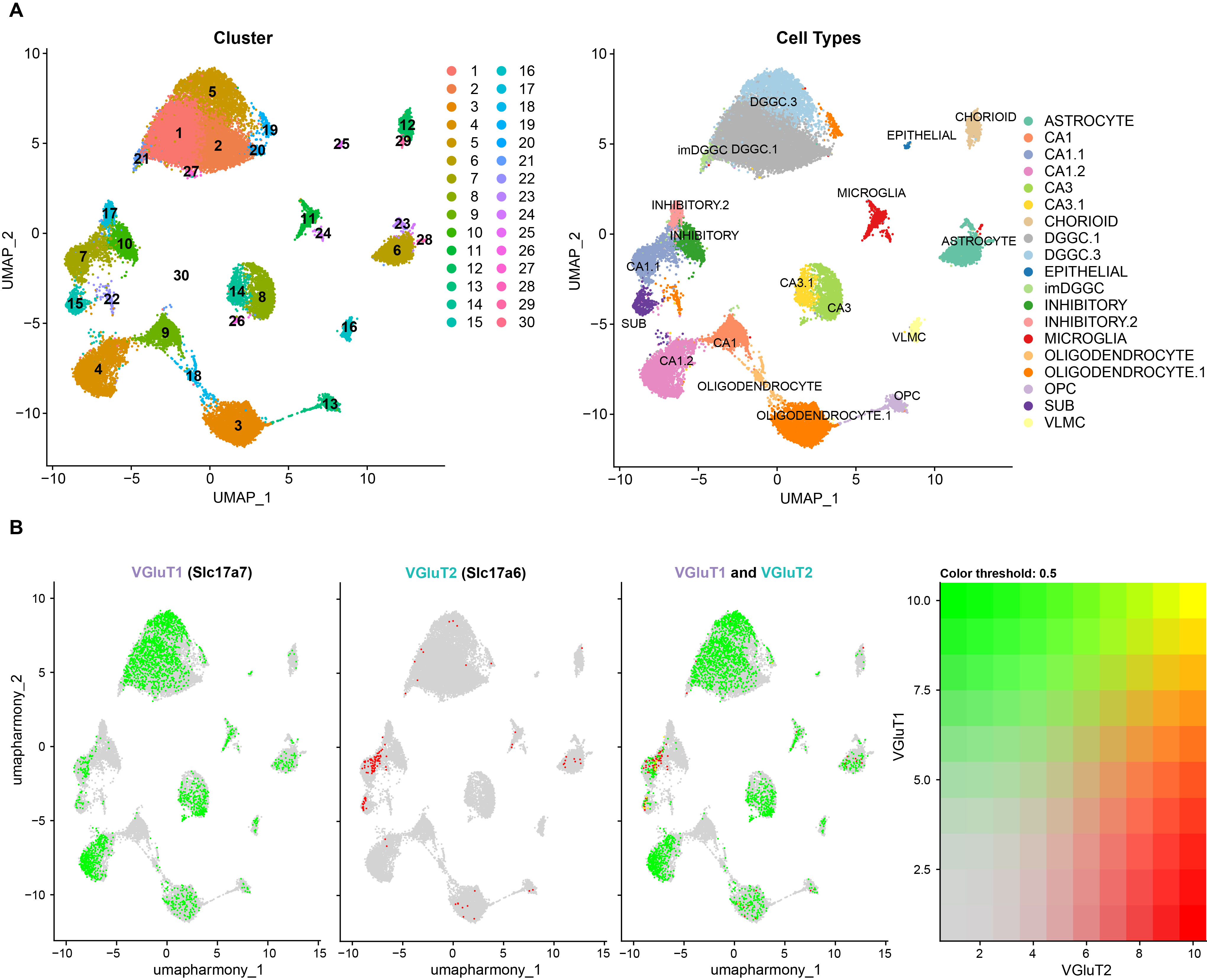

### Figure S5

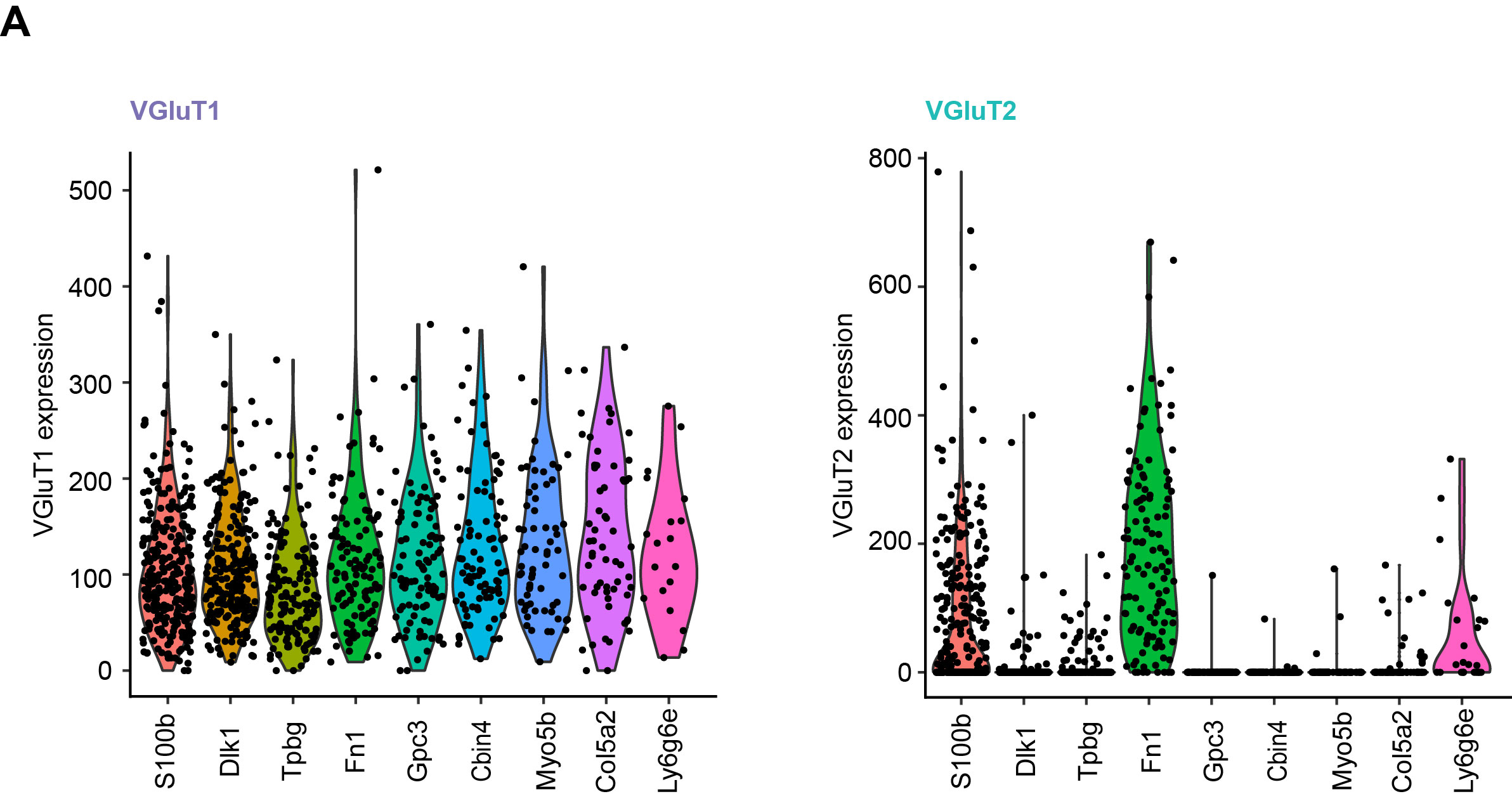

### Figure S6

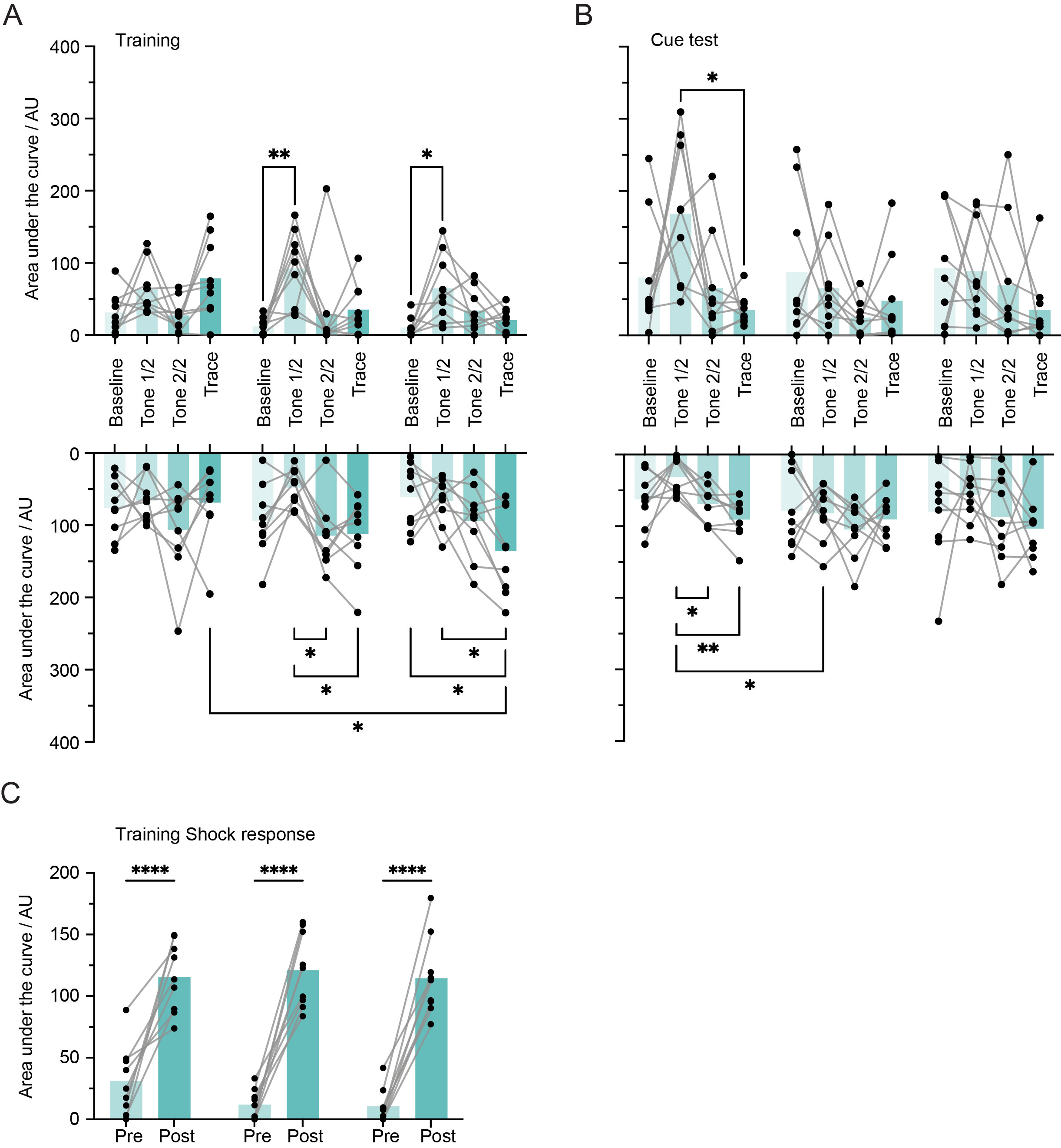

### Figure S7

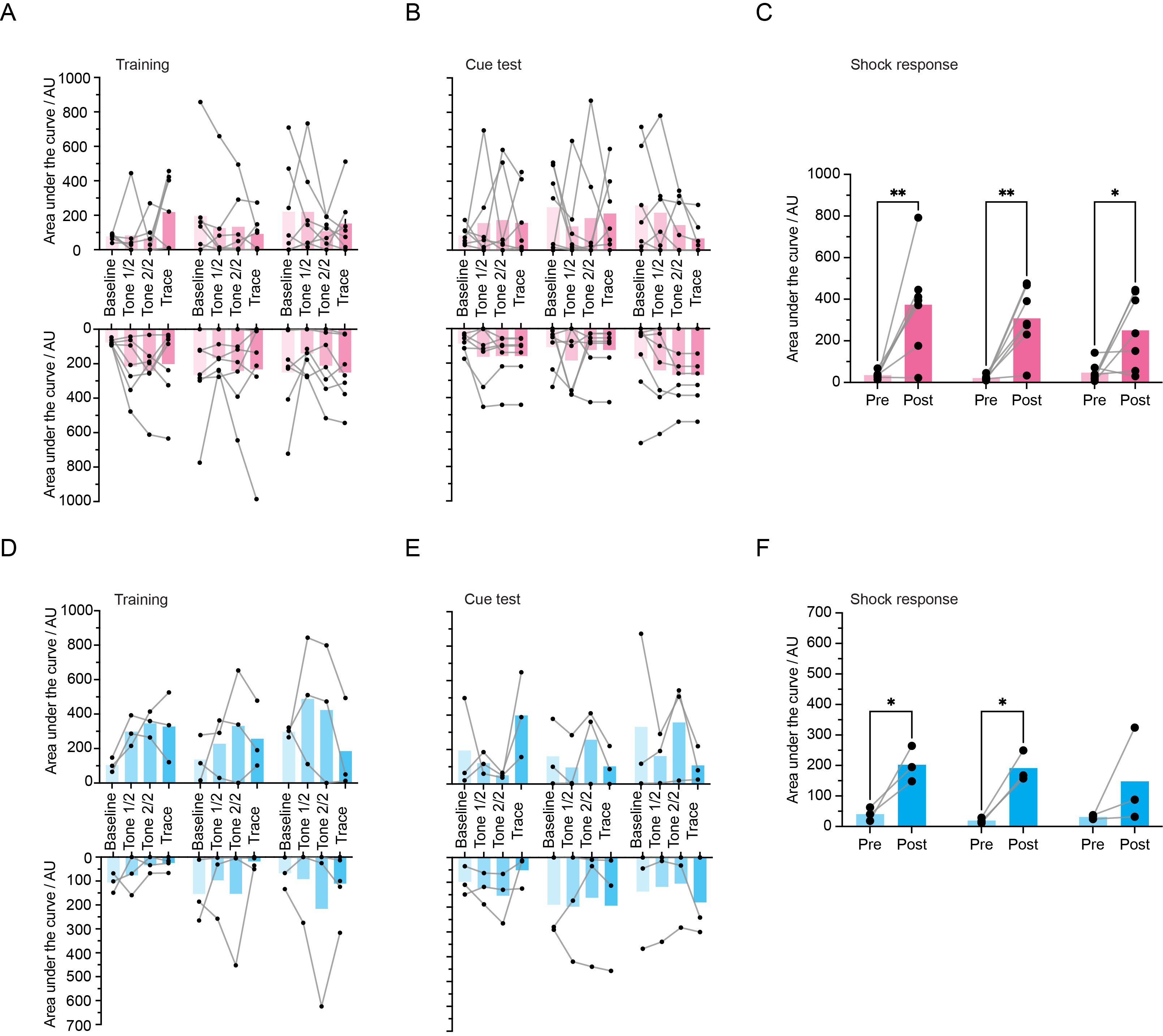

### Figure S8

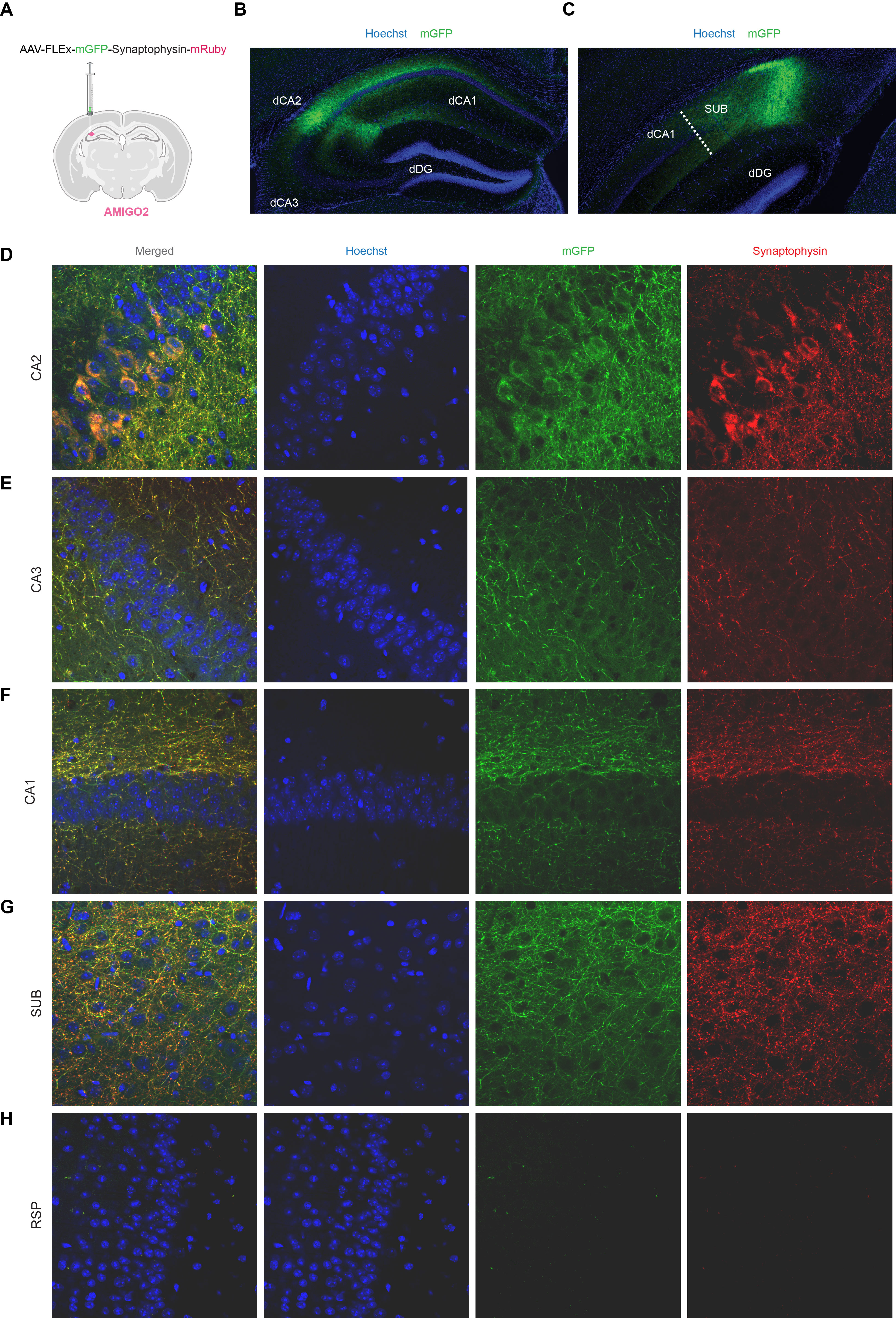

### Figure S9

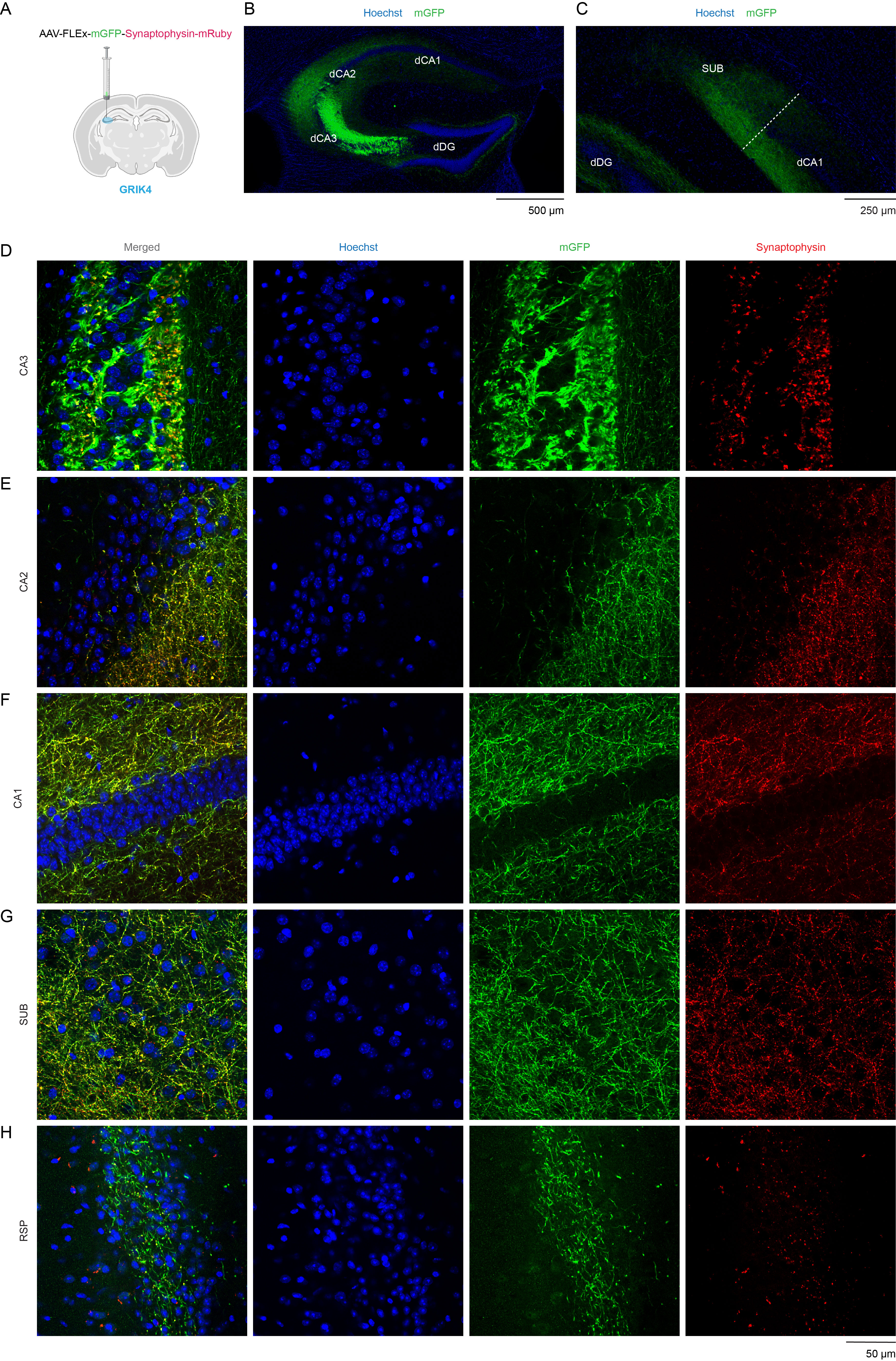
