## Supplementary figure legends for "Response dynamics of discrete subiculum→retrosplenial cortex projections underlying trace fear conditioning"

**Supplementary Figure 1. Role that VGluT1^+^ and VGluT2^+^ SUB→RSP afferents in TFC**

**A** Experimental diagram detailing AAV injections in the DH and canula placement in the RSP. **B** Left, Expression diagram detailing the locations of VGluT1 and VGluT2 positive DH afferents in the RSP. Right, representative microscopy images of the RSP in VGluT1 and VGluT2 subject mice. **C** Left, Experimental design of TFC behavior task, with CNO/Vehicle infusion taking place 30 minutes prior to the cue test. Middle, In male VGluT1-Cre mice, terminal silencing had no impact on freezing in response to tone or trace (n=8-7; two-way ANOVA with repeated measures; factor: treatment, P=0.6325, F_(1, 13)_= 0.2398, factor: phase, P<0.0001, F_(2, 26)_= 34.95, factor: treatment x phase, P=0.8653, F_(2, 26)_= 0.1455). Right, In male VGluT2-Cre mice, terminal silencing prior to testing significantly impaired freezing during the trace but not tone when compared to the vehicle group (n=7; two-way ANOVA with repeated measures; factor: treatment, P=0.0287, F_(1, 12)_= 6.174, factor: phase, P<0.0001, F_(2, 24)_= 48.11, factor: treatment x phase, P=0.0985, F_(2, 24)_= 2.557). **D** Left, Experimental design of used behavior task. The diagram depicts TFC, with CNO/Vehicle infusion taking prior to training in male mice, and a context test included on day 3. Middle, In male VGluT1-Cre mice, terminal silencing with CNO 30 minutes prior to training significantly impaired freezing response to context (n = 9; Unpaired two-tailed t-test; df=16, P=0.0064). Right, In male VGluT2-Cre mice, terminal silencing did not impaired freezing response to context (n = 10; Unpaired two-tailed t-test; df=18, P=0.2854). **E** Left, Experimental design of TFC behavior task conducted in female mice, with CNO/Vehicle infusion taking place 30 minutes prior to training. Middle, In female VGluT1-Cre mice, terminal silencing had no impact on freezing (n=7; two-way ANOVA with repeated measures; factor: treatment, P=0.5203, F_(1, 12)_= 0.4386, factor: phase, P<0.0001, F_(2, 24)_= 23.92, factor: treatment x phase, P=0.7125, F_(2, 24)_= 0.3438). Right, In female VGluT2-Cre mice, terminal silencing significantly impaired freezing during the trace but not tone compared to the vehicle group (n=6; two-way ANOVA with repeated measures; factor: treatment, P=0.0683, F_(1, 10)_= 4.173, factor: phase, P<0.0001, F_(2, 20)_= 21.32, factor: treatment x phase, P=0.0246, F_(2, 24)_= 4.483). **F** Left, Experimental design of used behavior task. The diagram depicts TFC, with CNO/Vehicle infusion taking prior to training in female mice, and a context test included on day 3. Middle, In female VGluT1-Cre mice, terminal silencing with CNO 30 minutes prior to training significantly impaired freezing response to context (n = 7; Unpaired two-tailed t-test, df=12; P=0.0205). Right, In female VGluT2-Cre mice, terminal silencing did not impaired freezing response to context (n = 6-5; Unpaired two-tailed t-test; df=9, P=0.1234).

**Supplementary Figure 2. Characterization of Delay, Trace, and Light Fear Conditioning**

**A** Left, Experimental design of TFC behavior task, with CNO/Vehicle infusion taking place 30 minutes prior to training. Middle, Experimental diagram detailing AAV injections in the DH and canula placement in the RSP. In this control experiment, subject mice receive a tracer AAV as opposed to an inhibitory DREADD AAV, to examine the effect of CNO administration on TFC. Right, CNO infusion 30 minutes prior to training had no effect on freezing behavior as compared to the vehicle control (n = 9-8; two-way ANOVA with repeated measures; factor: treatment, P=0.4931, F_(1, 15)_= 0.4935, factor: phase, P<0.0001, F_(2, 30)_= 30.53, factor: treatment x phase, P=0.9253, F_(2, 30)_= 0.07789). **B** Left, Experimental design of paired TFC and DFC behavior task. Right, DFC animals exhibited significantly less freezing during trace, but not tone, as compared to TFC group (n = 8-9; two-way ANOVA with repeated measures; factor: paradigm, P=0.0035, F_(1, 15)_= 12.01, factor: phase, P<0.0001, F_(2, 30)_= 24.49, factor: paradigm x phase, P=0.0020, F_(2, 30)_= 7.667). **C** Left, Experimental design of paired TFC and TLC behavior task. Top Right, TFC animals exhibited significantly higher freezing during the trace period on day 2, as compared against animals that underwent TLC (n = 7-6; two-way ANOVA with repeated measures; factor: paradigm, P=0.0572, F_(1, 11)_= 4.511, factor: phase, P<0.0001, F_(2, 22)_= 41.39, factor: paradigm x phase, P=0.0081, F_(2, 22)_= 6.046). Bottom Right, TLC animals exhibited significantly higher freezing during light presentation on day 3, as compared against animals that underwent TFC (n = 7-6; two-way ANOVA with repeated measures; factor: paradigm, P=0.0115, F_(1, 11)_= 9.164, factor: phase, P<0.0001, F_(2, 22)_= 36.99, factor: paradigm x phase, P=0.0356, F_(2, 22)_= 3.898).

**Supplementary Figure 3. Proteomic characterization of VGluT1^+^ and VGluT2^+^ SUB→RSP projections**

**A** Experimental workflow for *in vivo* proximity biotinylation assay of SUB VGluT1^+^ and VGluT2^+^ projections in the RSP. **B** KEGG analysis showing the differences between vGluT1- and vGluT2-containing presynaptic functional proteomes. **C** The levels of biotinylated cytosolic proteins in VGluT1^+^ and VGluT2^+^ RSP terminals were similar, as indicated by significant correlation.

**Supplementary Figure 4. Characterization of VGluT1 and VGlut2 expression amongst dorsal hippocampus cell types**

**A** Left, Raw single cell RNA Seq clustering of neuronal and non-neuronal cell types within the dorsal hippocampus using Uniform Manifold Approximation and Projection (UMAP). Right, Clustered cell populations with their corresponding cell type identities using known neuronal and non-neuronal cell markers. **B** Far Left & Middle Left, VGluT1 and VGluT2 expression patterns within clustered cell types. VGluT1 is expressed widely across numerous dorsal hippocampal cell types, whereas VGluT2 is mainly expressed in CA1 and subiculum cell types. Middle Right, VGluT1 and VGluT2 expression overlap amongst clustered cell types. Far Right, VGluT1 and VGluT2 expression legend.

**Supplementary Figure 5. Cluster-specific expression of VGluT1 and VGlut2 in the SUB**

**A** Violin plots of VGluT1 and VGlut2 expression across SUB clusters, identified by cluster-specific marker genes. VGlut2 shows more conserved expression predominantly in S100b, Fn1, and Ly6g6e clusters, compared to VGlut1 which is expressed across all clusters.

**Supplementary Figure 6. Area Under the Curve quantification for VGluT2^+^ SUB→RSP afferents**

**A** Change in GCaMP area-under-the-curve (AUC) between phases at training day (n=9; two-way ANOVA with repeated measures; Positive Peak, top, factor: trial, P=0.1689, F_(1.790, 57.27)_= 1.859, factor: phase, P<0.0001, F_(3, 32)_= 11.88, factor: trial x phase, P= 0.0812, F_(6, 64)_= 1.982; Negative Peak, bottom, factor: trial, P=0.4893, F_(1.869, 59.80)_= 0.7044, factor: phase, P=0.0008, F_(3, 32)_= 7.242, factor: trial x phase, P=0.0955, F_(6, 64)_= 1.894). **B** Change in GCaMP area-under-the-curve (AUC) between phases at testing day (n=9; two-way ANOVA with repeated measures; Positive Peak, top, factor: trial, P=0.1995, F_(1.872, 59.92)_= 1.664, factor: phase, P=0.0044, F_(3, 32)_= 5.303, factor: trial x phase, P=0.1726, F_(6, 64)_= 1.563; Negative Peak, bottom, factor: trial, P=0.0453, F_(1.731, 55.40)_ = 3.444, factor: phase, P=0.0194, F_(3, 32)_= 3.804, factor: trial x phase, P=0.6614, F_(6, 64)_= 0.6862). **C** Increase in average AUC following shock exposure (n=9; two-way ANOVA with repeated measures; factor: trial, P=0.3864, F_(1.894, 30.30)_= 0.9699, factor: phase, P<0.0001, F_(1, 16)_= 164.4, factor: trial x phase, P=0.2625, F_(2, 32)_= 1.395).

**Supplementary Figure 7. Area Under the Curve quantification for CA2 and CA3 pyramidal cell bodies**

**A** No significant changes were observed in GCaMP area-under-the-curve (AUC) between phases at training (n=7; two-way ANOVA with repeated measures; Positive Peak, top, factor: trial, P=0.3812, F_(2, 48)_= 0.9840, factor: phase, P=0.8345, F_(3, 24)_= 0.2868, factor: trial x phase, P=0.4807, F_(6, 48)_= 0.9322; Negative Peak, bottom,  factor: trial, P=0.5697, F_(2, 48)_= 0.5694, factor: phase, P=0.8939, F_(3, 24)_= 0.2022, factor: trial x phase, P=0.3547, F_(6, 48)_= 1.139). **B** No significant changes were observed in GCaMP area-under-the-curve (AUC) between phases at testing (n=7; two-way ANOVA with repeated measures; Positive Peak, top, factor: trial, P=0.6824, F_(2, 48)_= 0.3852, factor: phase, P=0.9180, F_(3, 24)_= 0.1664, factor: trial x phase, P=0.6817, F_(6, 48)_= 0.6605; Negative Peak, bottom, factor: trial, P=0.0222, F_(2, 48)_= 4.127, factor: phase, P=0.6005, F_(3, 24)_= 0.6337, factor: trial x phase, P=0.9783, F_(6, 48)_= 0.1896). **C** Repeated significant increase in average AUC following shock exposure (left, n=7; two-way ANOVA with repeated measures; factor: trial, P=0.4501, F_(1.541, 18.49)_= 0.7581, factor: phase, P=0.0002, F_(1, 12)_= 29.01, factor: trial x phase, P=0.3541, F_(2, 24)_= 1.085). **D** No significant changes were found in GCaMP signal area-under-the-curve (AUC) between phases at training (n=3; two-way ANOVA with repeated measures; Positive Peak, top, factor: trial, P=0.3165, F_(2, 16)_=1.2370, factor: phase, P=0.5742, F_(3, 16)_= 0.7072, factor: trial x phase, P=0.6254, F_(6, 16)_=0.7400; Negative Peak, bottom, factor: trial, P=0.2718, F_(2, 20)_= 1.3910, factor: phase, P=0.7855, F_(3, 10)_= 0.3568, factor: trial x phase, P=0.4297, F_(6, 20)_=1.0390). **E** No significant changes were found in GCaMP signal area-under-the-curve (AUC) between phases at testing, except for significant trial x phase interaction for positive peak (right, n=3; two-way ANOVA with repeated measures; Positive Peak, top, factor: trial, P=0.3508, F_(2, 16)_= 1.1190, factor: phase, P=0.9075, F_(3, 8)_= 0.1793, factor: trial x phase, P=0.0483, F_(6, 16)_=2.771; Negative Peak, bottom, factor: trial, P=0.2069, F_(2, 16)_=1.7420, factor: phase, P>0.9999, F_(3, 8)_= 0.0011, factor: trial x phase, P=0.8430, F_(6, 16)_= 0.4377). **F** Increase in average AUC following shock exposure reduces with repeated trials (left, n=3; two-way ANOVA with repeated measures; factor: trial, P=0.7141, F_(1.241, 4.966)_= 0.2135, factor: phase, P=0.0007, F_(1, 4)_= 87.91, factor: trial x phase, P=0.8335, F_(2, 8)_= 0.1863).

**Supplementary Figure 8. Characterization of CA2 connectivity**

**A** Diagram depicting a unilateral CA2 injection of a *cre*-dependent synaptophysin pAAV1-hSyn-FLEx-mGFP-2A-Synaptophysin-mRuby viral vector in Amigo2-cre mice. **B** Fluorescence microscopy image showing viral expression in the dorsal CA2 and surrounding hippocampal subfields. **C** Fluorescence microscopy image of the dorsal SUB, showing direct projection from the CA2 into the SUB. **D-H** Confocal microscopy images depicting GFP (green) and synaptophysin (red) expression in various hippocampal and extrahippocampal brain regions. The dorsal CA2 pyramidal cells form synaptic connections in the CA3, CA1, and SUB, but not in the RSP.

**Supplementary Figure 9. Characterization of CA3 connectivity**

**A** Diagram depicting a unilateral CA3 injection of a *cre*-dependent synaptophysin pAAV1-hSyn-FLEx-mGFP-2A-Synaptophysin-mRuby viral vector in Grik4-Cre mice. **B** Fluorescence microscopy image showing viral expression in the dorsal CA3 and surrounding hippocampal subfields. **C** Fluorescence microscopy image of the dorsal SUB, showing direct projection from the CA3 into the subiculum. **D-H** Confocal microscopy images depicting GFP (green) and synaptophysin (red) expression in various hippocampal and extrahippocampal brain regions. The dorsal CA3 pyramidal cells form synaptic connections in the CA1, CA2, CA3, SUB, and RSP.
